# The SARS-CoV-2 spike protein binds and modulates estrogen receptors

**DOI:** 10.1101/2022.05.21.492920

**Authors:** Oscar Solis, Andrea R. Beccari, Daniela Iaconis, Carmine Talarico, Camilo A. Ruiz-Bedoya, Jerome C. Nwachukwu, Annamaria Cimini, Vanessa Castelli, Riccardo Bertini, Monica Montopoli, Veronica Cocetta, Stefano Borocci, Ingrid G. Prandi, Kelly Flavahan, Melissa Bahr, Anna Napiorkowski, Giovanni Chillemi, Masato Ooka, Xiaoping Yang, Shiliang Zhang, Menghang Xia, Wei Zheng, Jordi Bonaventura, Martin G. Pomper, Jody E. Hooper, Marisela Morales, Avi Z. Rosenberg, Kendall W. Nettles, Sanjay K. Jain, Marcello Allegretti, Michael Michaelides

**Affiliations:** Biobehavioral Imaging and Molecular Neuropsychopharmacology Unit, National Institute on Drug Abuse Intramural Research Program, Baltimore, 21224, MD, USA; EXSCALATE, Dompé farmaceutici S.p.A, Napoli, Italy; Center for Infection and Inflammation Imaging Research, Johns Hopkins University School of Medicine, Baltimore, MD, USA; Department of Pediatrics, Johns Hopkins University School of Medicine, 1550 Orleans Street, CRB-II Room 109, Baltimore, MD, USA; Center for Tuberculosis Research, Johns Hopkins University School of Medicine, Baltimore, MD, USA; Department of Integrative Structural and Computational Biology, The Scripps Research Institute, 130 Scripps Way, Jupiter, FL 33458, USA; Department of Life, Health and Environmental Sciences, University of L’Aquila, L’Aquila, Italy; Sbarro Institute for Cancer Research and Molecular Medicine, Department of Biology, Temple University, Philadelphia, PA, USA; Atreius S.a.s., L’Aquila, Italy; Department of Pharmaceutical and Pharmacological Sciences, University of Padova, Padova, Italy; VIMM- Veneto Institute of Molecular Medicine, Fondazione per la Ricerca Biomedica Avanzata, Padova, Italy; Department for Innovation in Biological, Agro-Food and Forest Systems, DIBAF, University of Tuscia, Viterbo, Italy; Division of Preclinical Innovation, National Center for Advancing Translational Sciences, Rockville, MD, USA; Department of Pathology, Johns Hopkins University School of Medicine, Baltimore, MD, USA; Neuronal Networks Section, National Institute on Drug Abuse Intramural Research Program, Baltimore, 21224, MD, USA; Departament de Patologia i Terapèutica Experimental, Institut de Neurociències, Universitat de Barcelona, L’Hospitalet de Llobregat, Catalonia; Department of Radiology and Radiological Sciences, Johns Hopkins University School of Medicine, Baltimore, MD, USA; Department of Pathology, Stanford University School of Medicine, Stanford, CA, USA; Dompé farmaceutici S.p.A, L’Aquila, Italy; Department of Psychiatry & Behavioral Sciences, Johns Hopkins School of Medicine, Baltimore, MD, USA

**Author notes:** Corresponding authors: Michael Michaelides, Ph.D., National Institute on Drug Abuse, 251 Bayview Blvd, Baltimore, MD 21224, Marcello Allegretti, ChemD., Dompé farmaceutici S.p.A, Via Campo di Pile s.n.c. - 67100 L’Aquila, Italy.

## Abstract

The severe acute respiratory syndrome coronavirus 2 (SARS-CoV-2) spike (S) protein binds angiotensin-converting enzyme 2 (ACE2) at the cell surface, which constitutes the primary mechanism driving SARS-CoV-2 infection. Molecular interactions between the transduced S and endogenous proteins likely occur post-infection, but such interactions are not well understood. We used an unbiased primary screen to profile the binding of full-length S against >9,000 human proteins and found significant S-host protein interactions, including one between S and human estrogen receptor alpha (ERα). After confirming this interaction in a secondary assay, we used bioinformatics, supercomputing, and experimental assays to identify a highly conserved and functional nuclear receptor coregulator (NRC) LXD-like motif on the S2 subunit and an S-ERα binding mode. In cultured cells, S DNA transfection increased ERα cytoplasmic accumulation, and S treatment induced ER-dependent biological effects and ACE2 expression. Noninvasive multimodal PET/CT imaging in SARS-CoV-2-infected hamsters using [^18^F]fluoroestradiol (FES) localized lung pathology with increased ERα lung levels. Postmortem experiments in lung tissues from SARS-CoV-2-infected hamsters and humans confirmed an increase in cytoplasmic ERα expression and its colocalization with S protein in alveolar macrophages. These findings describe the discovery and characterization of a novel S-ERα interaction, imply a role for S as an NRC, and are poised to advance knowledge of SARS-CoV-2 biology, COVID-19 pathology, and mechanisms of sex differences in the pathology of infectious disease.

## Main Text

COVID-19 is an infectious disease caused by the severe acute respiratory syndrome coronavirus 2 (SARS-CoV-2). The most frequent symptom of severe COVID-19 is pneumonia, accompanied by fever, cough, and dyspnea commonly associated with cytokine storm, systemic inflammatory response, and coagulopathy^1,2^. The elderly and those with underlying comorbidities, are more likely to develop severe illness and mortality^3,4^.

SARS-CoV-2 is characterized by four structural proteins: spike (S), envelope (E), membrane (M), and nucleocapsid (N) proteins^5^. Currently, most COVID-19 vaccines use S as the target antigen, as it is an important determinant capable of inducing a robust protective immune response^6^. Furthermore, it is a critical component for cell infection via direct interaction with ACE2^7,8^. S is composed of 1273 amino acids. It consists of a signal peptide located at the N-terminus (amino acids 1–13), the S1 subunit (14–685 residues) and the S2 subunit (686–1273 residues). The S1 subunit contains the ACE2 receptor-binding domain (RBD), whereas the S2 subunit is responsible for viral and host cell membrane fusion^9,10^ which requires other proteins^8,11-14^. Importantly, cells are still susceptible to infection and show S-dependent biological responses independent of ACE2^15–19^ suggesting that S may promote pathology independent from its capacity to bind ACE2. Given the multitude and complex array of systemic symptoms associated with COVID-19, it is possible that other molecular targets of S may exist. Identification of additional S targets would be critical for advancing our understanding of SARS-CoV-2 infection and COVID-19 pathobiology.

### S binds ERα with high affinity

First, we radiolabeled full-length recombinant S with ^125^I and confirmed its binding to human ACE2 (K_D_=27.8 ± 5.0 nM)^13,20^ (**Fig. 1a, b**). Next, we used Protoarray®protein array slides to screen for [^125^I]S binding against >9000 human proteins **(Fig. 1c)**. As per Protoarray®protocol, we also tested the binding of [^3^H]estradiol (E2) (1 nM), which serves as a positive control. As expected, incubation with [^3^H]E2 labeled the full-length estrogen receptor alpha (ERα) (**Fig. 1d, f**). Other arrays were incubated with [^125^I]S (20 nM) with or without [^3^H]E2 (1 nM) or in the presence of non-radioactive S (300 nM) to control for non-specific (NS) binding (**Fig. 1e**). We detected a specific [^125^I]S signal at seven proteins on the array, including neuropilin 1 (NRP1) (**Extended Table 1**), a known S target protein^12^. Surprisingly, we also detected a specific and reproducible [^125^I]S signal at the multiple ERα sites (**Fig. 1g-i**). We then confirmed the S-NRP1 and S-ERα interactions in secondary assays by immobilizing S and performing surface plasmon resonance (SPR) kinetic analyses with recombinant ACE2 (K_D_= 0.58 nM), NRP1 (K_D_= 89.4 nM), and ERα (K_D_ = 9.7 nM) (**Fig. 1j-l**).

**Figure 1.**
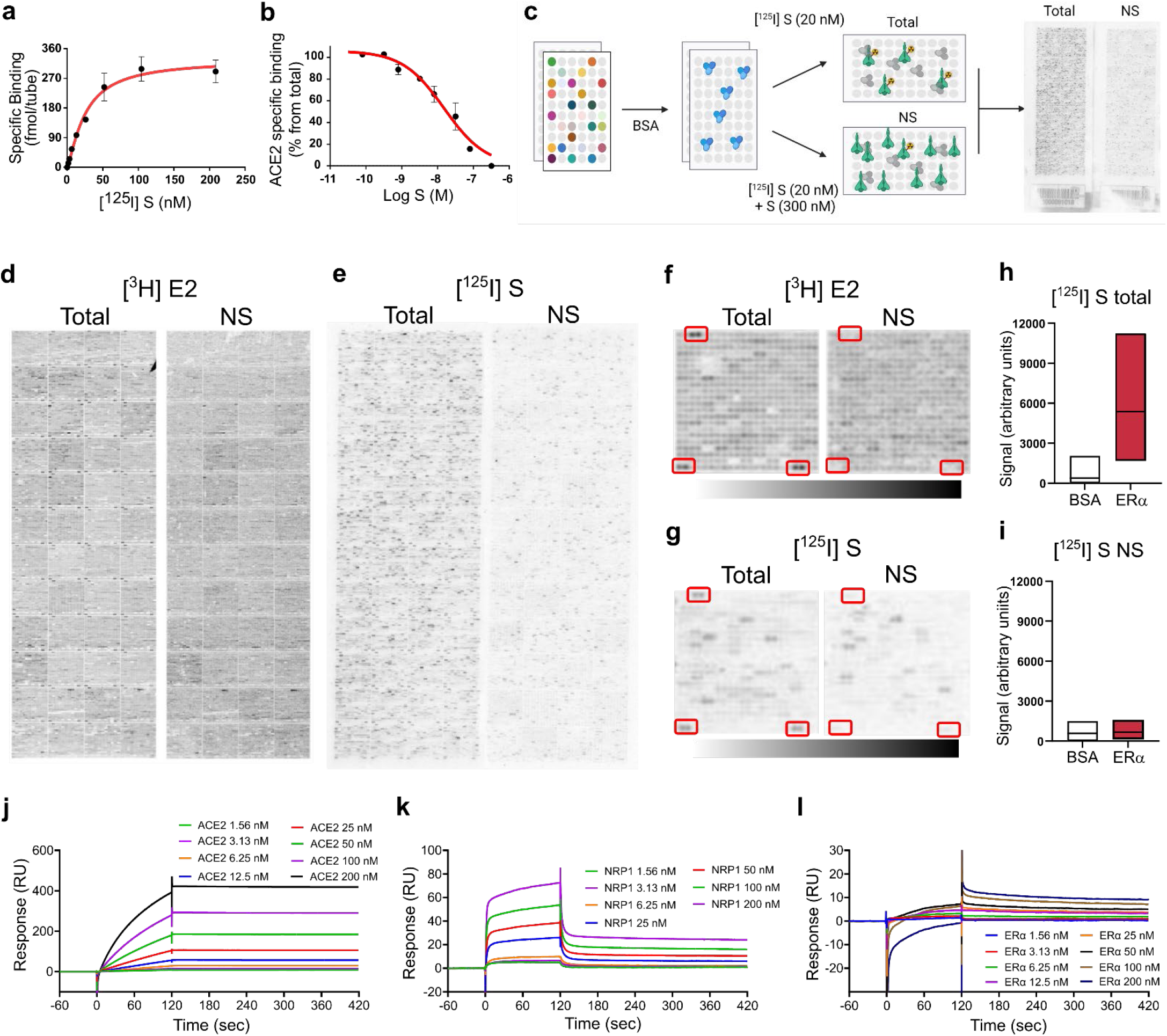
S binds ERα with high affinity. (**a**) [^125^I]S saturation and (**b**) competition binding to recombinant ACE2. (**c**) Schematic of ProtoArray^®^ experimental design. (**d**) Positive control ProtoArray^®^ autoradiograms showing total and non-specific (NS) binding of [^3^H]estradiol (E2). (**e**) ProtoArray^®^ autoradiograms showing total and NS binding of [^125^I]S. (**f, g**) Representative array blocks showing total and NS [^3^H]E2 and [^125^I]S binding. Red rectangles show location of ERα proteins. (**h, i**) Quantification of total and NS [^125^I]S binding at ERα and BSA (control). Data are representative of three independent experiments. (**j-l**) Representative SPR sensorgrams showing kinetic and equilibrium binding analyses of immobilized S exposed to increasing concentrations of ACE2, NRP1 and ERα protein (K_on_=2.03 x10^5^, K_off_= 1.96 x 10^-3^, K_D_= 9.7 nM). In a–b, data are represented as mean ± SEM. In h, i, data are presented as median ± min and max limits.

### S and ER interact at conserved LXD-like nuclear receptor coregulator (NRC) motifs

To identify discrete structural domains involved in S-ER interactions we used bioinformatics and the EXaSCale smArt pLatform Against paThogEns (EXSCALATE) supercomputing platform^21^. First, a network analysis confirmed prominent interactions between ERα, ERβ, and other proteins (**Fig. 2a**) including known interactions with NR coactivators 1, 2, and 3 (NCOA1, NCOA2, NCOA3) (**Extended Table 2**). NCOAs bind to the activation function 2 (AF-2) region on ERs to modulate ligand-mediated activation of ER transcription via a region called the NR box which includes an LXD motif, known as the LXXLL core consensus sequence (where L is leucine and X is any amino acid) ^22^ (**Extended Data Fig. 1**). This motif is necessary and sufficient for NRC binding to ligand-bound ERs and for ER function. Using this information and the EXSCALATE platform, we identified two ER-interacting LXD-like motifs in the S sequence (**Fig. 2b**). We then analyzed and compared the sequences of other coronavirus S proteins, including SARS-CoV, MERS-CoV (Middle East Respiratory Syndrome coronavirus), HCoV (Human coronavirus) and MHV (Murine coronavirus) (**Extended Table 3**) to search for conserved LDX-like motifs. We found discrete shared amino acid patterns across species (**Extended Data Fig. 2**), suggesting a conserved functional role of these regions. We then verified the conservation of the two LDX-like motifs and found the LPPLL pattern at residues 861-865 conserved among SARS-CoV-2, SARS-CoV, HCoV and MERS-CoV (**Fig. 2c, Extended Data Fig. 2**), while the LXD-like pattern IEDLL at residues 818-822 is also conserved among the same viruses. It is also worth noting that a standard LEDLL pattern is found in HCoV-HKU1 in the same position. Notably, this LXD-like region, which is solvent-exposed in the S experimental structures, retains well-defined 3D structural characteristics (alpha-helix folding, red in **Fig. 2c**) found in the ER-NCOA complexes 3UUD^23^ and 3OLL^24^. On the contrary, the LXXLL motif, less solvent-exposed, is unstructured (blue in **Fig. 2c**). It is well known, however, that the motif region may assume the alpha-helix folding only after the binding with ER^25^, implying a conformational rearrangement of the two molecular partners.

**Figure 2.**
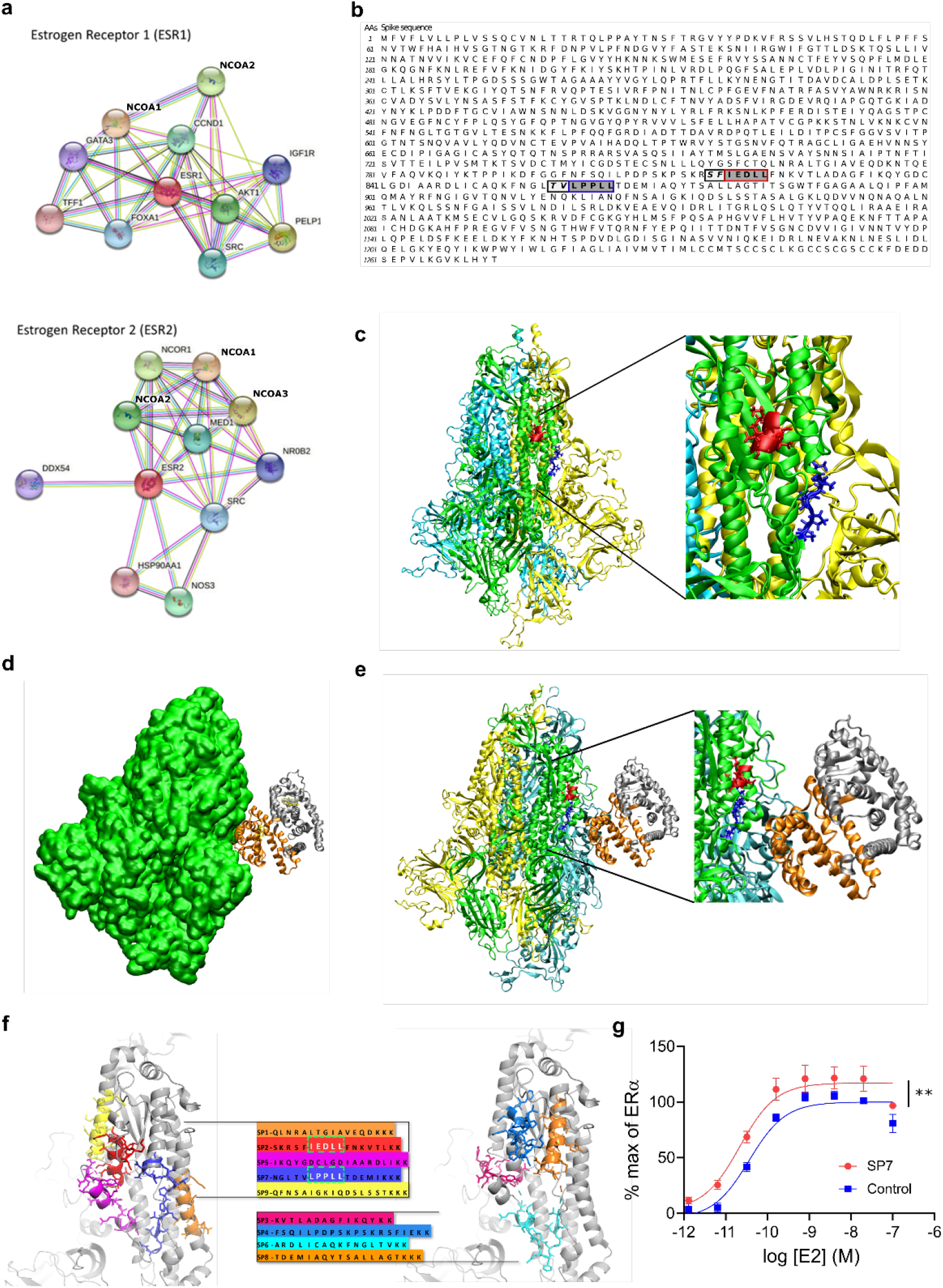
S and ER interact at conserved LXD nuclear receptor coregulatory (NRC) motifs. (**a**) ER interaction network showing known and predicted protein associations. (**b**) LXD-like patterns in the S sequence. The LXXLL motif and a homologous region are highlighted in blue and red boxes, respectively, with dark grey background. −1 and −2 positions are reported in italic and light grey background. (**c**) The LPPLL and IEDLL residues of the two motifs are shown in the 3D X-ray S structure (pdb id 6VYB) with blue and red colors, respectively. The three S chains are shown in yellow, cyan and green. (**d**) S-ER motif-oriented docking. The best 3D docking hypothesis is shown. The ER dimer is in orange and gray, S is green. (**e**) Alignment between the best-pose and the 3OLL model tied with NCOA1. The region occupied by S’s alpha-helix interacts in the area where the NCOA fragment was crystallized. (**f**) S protein peptides and their location with respect to the S 3D structure. (**g**) The SP7 peptide containing the LPPLL motif significantly increased ERα activation (F (1,48) = 30.38; **P<0.01; two-way ANOVA, peptide treatment main effect). Data are mean ± SEM.

We then performed *in silico* molecular docking simulations to identify a putative S-ER binding mode. An S-ERα 3D model was built based on PDB 6VYB in its wild-type and fully glycosylated form and PDB 3OLL, which contained both E2 and NCOA1^24^. For protein-protein docking, S in glycosylated form and ERα/β were used as receptor and ligand, respectively. The top 100 predicted complex structures were selected, and the ten best hypotheses were visually inspected to confirm the reliability of the calculation^26^. Since the revised LXD-like motifs identified were located outside the S RBD, the ability of ER to interact outside this region was evaluated by means of blind-docking^26^. The best binding hypothesis included evidence of a high-affinity ER interaction towards the lateral region of S, which includes the so-called “fusion peptide portion” (**Extended Data Fig. 3**). The structural information that ER residues are recognized by NCOA was then used to guide S-ER docking studies by optimizing protein-protein interactions. The best binding hypothesis obtained highlighted the binding of ER to the S region containing the two described LDX-like motifs (**Fig. 2d, e**). Several Molecular Dynamics (MD) simulations of the best docking complexes were then carried out. MD results showed the formation of a strong interaction between ER and S even in the first phase of the recognition (**Extended Data Fig. 4**) We then extracted 9 peptide sequences (SP1-9) based on their proximity to the putative S-ER binding region and their LXD-like domain sequence similarities (**Fig. 2f**), synthesized each peptide and examined their effects on ERα-mediated transcriptional activation (GeneBLAzer™ ERα/β-UAS-bla GripTite™ cells). One peptide, (SP7), which solely contained the LPPLL motif, significantly increased the potency of E2 in stimulating ERα transcriptional activation (**Fig. 2g, Extended Data Fig. 5**).

### S modulates ER-dependent biological functions

We used MCF-7 nuclear extracts and the TransAM™ ER assay^27^ to measure E2-stimulated ERα DNA binding. We found that full-length S (IC_50_ = 2.4 ± 1.5 nM) and S trimer (IC_50_ = 72 ± 2.6 nM), but not S-RBD, inhibited E2-stimulated ERα DNA binding (**Fig. 3a, Extended Data Fig. 6a, b**). We also assessed whether S affected ER-mediated transcriptional activation using ERα-LBD and ERβ-LBD reporter cell lines (GeneBLAzer™ ERα/β-UAS-bla GripTite™). S overexpression significantly decreased the E_max_ of E2-stimulated ERα (E_max_^S^ = 72 ± 8.8%; E_max_^Control^ = 100%) and ERβ (E_max_^S^ = 85 ± 2.3 %; E_max_^Control^ = 100%) transcriptional activation without significantly affecting their EC_50_ (ERα: EC_50_^S^: 0.15 ± 0.04 nM; EC_50_^Control^: 0.23 ± 0.05 nM and ERβ: EC_50_^S^: 0.51 ± 0.03 nM; EC_50_^Control^: 0.70 ± 0.06 nM) (**Fig. 3b, Extended Data Fig. 6c**) suggesting a selective partial antagonism of the ER-induced transcriptional effect.

**Figure 3.**
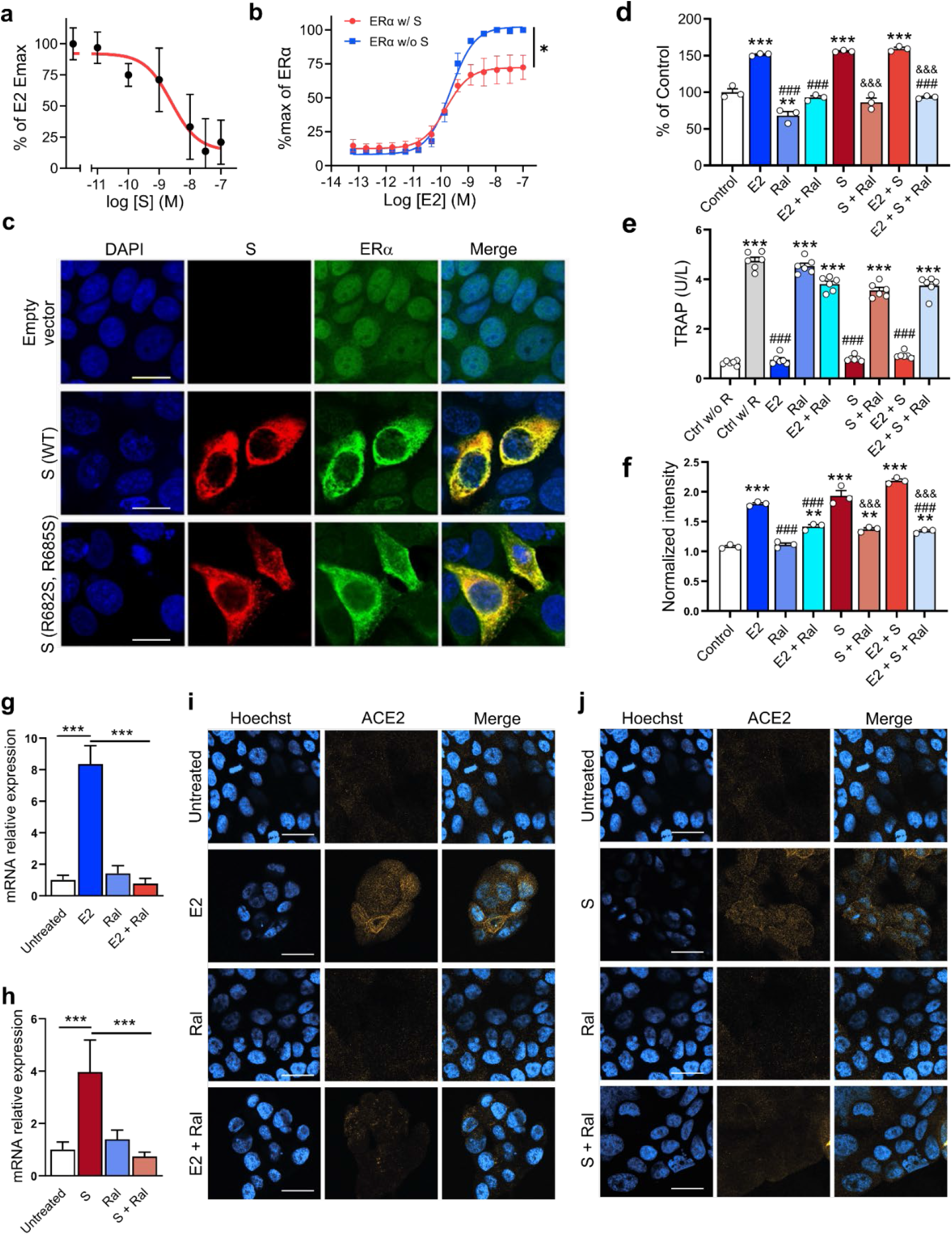
S modulates ER-dependent biological functions. S inhibits (**a**) E2-induced ERα DNA binding in MCF-7 nuclear extracts and (**b**) transcriptional activation in an ERα reporter cell line (F (1, 28) = 21.73, \**P*=0.01; two-way ANOVA, S treatment x E2 concentration interaction effect). (**c**) Immunofluorescent staining of S and endogenous ERα in MCF-7 cells transfected with empty vector, wild-type (WT) or the furin cleavage site mutant S^(R682S, R685S)^. Scale bar = 16 μm. (**d**) S increases MCF-7 cell proliferation in an ER-dependent manner (\*\**P*<0.01, \*\*\**P*<0.001 versus control; ^###^*P*<0.001 versus E2; ^&&&^*P*<0.001 versus S; one-way ANOVA with post hoc Tukey test). (**e**) S decreases osteoclast differentiation in an ER-dependent manner (***p<0.001 versus control w/o RANKL; ^###^*P*<0.001 versus control w/ RANKL; one-way ANOVA with post hoc Tukey test). (**f**) S and E2 increase ACE2 protein levels in MCF-7 cells in an ER-dependent manner (\*\*\**P*<0.001 versus control; ^###^*P*<0.001 versus E2; ^&&&^*P*<0.001 versus S, one-way ANOVA with post hoc Tukey test). (**g, h**) E2 and S increase ACE2 mRNA (\*\*\**P*<0.001, one-way ANOVA with post hoc Tukey test) and (**i, j**) protein in the Calu-3 lung cell line in an ER-dependent manner. Scale bar = 30 μm. All data shown as mean ±SEM.

To visualize the cellular distribution of S and ERα we transfected MCF-7 cells with the wild-type (WT) S or a mutant S^(R682S, R685S)^ stabilized at the furin cleavage site and performed immunocytochemistry. Cells transfected with the empty pcDNA3.1 vector showed the expected ERα nuclear-enriched distribution pattern and no S signal (**Fig. 3c**). In contrast, overexpression of either WT or mutant S increased ERα cytoplasmic labeling (**Fig. 3c**), indicating that S, either with or without an intact furin cleavage site, leads to an increase in ERα and its redistribution from the nucleus to the cytoplasm. Notably, the S-induced increase in cytoplasmic ERα labeling was not due to increases in ERα mRNA, though S transfection did significantly alter the expression of GREB1, a known ERα-target gene (**Extended Data Fig. 7**).

E2 increases MCF-7 cell proliferation, whereas raloxifene, a potent selective ER modulator (SERM), blocks MCF-7 cell proliferation^28^. As expected, E2 treatment increased MCF-7 cell proliferation and this effect was blocked by raloxifene (2 μM) (**Fig. 3d**). Intriguingly, S (10 ng/ml) itself also increased MCF-7 cell proliferation, and this effect was also blocked by raloxifene (2 μM) indicating it was ER-dependent. Notably, exposure of MCF-7 cells to E2 and S did not lead to an additive proliferation response and neither E2 nor S induced proliferation in an ER-lacking cell line (MDA-MB-231) (**Extended Data Fig. 8**).

E2 inhibition of osteoclast differentiation is an ERα-dependent effect linked to its therapeutic use^21,29,30^. RAW264.7, a murine macrophage cell line that expresses ERα, was induced to differentiate into osteoclasts by receptor activator of NF-κB ligand (RANKL) treatment in the presence or absence of either E2 (1 nM), S (10 ng/ml), or their combination. E2 or S, as well as their combination, abolished RANKL-induced osteoclast differentiation (**Fig. 3e**) and these effects were completely blocked by raloxifene (2 μM), indicating they were ER-dependent.

To assess the relevance of S and ER signaling to SARS-CoV-2 cell entry mechanisms, we first assessed the effect of E2 (1 nM) and S (10 ng/ml) on ACE2 levels in MCF-7 cells via ELISA. E2 or S, as well as their combination, significantly increased ACE2 levels and in both cases, these effects were blocked by raloxifene (2 μM) (**Fig. 3f**), indicating the ACE2 increases were ER-dependent. We also tested the effect of S and E2 on ACE2 expression in Calu-3 cells, a human airway epithelial cell line used to study SARS-CoV-2 infection. Both E2 (200 nM) and S (10 ng/mL) increased ACE2 mRNA (**Fig. 3g, h**) and ACE2 membrane protein expression (**Fig. 3i, j**). In both cases, raloxifene (20 μM) reverted these effects indicating they were ER-dependent.

### SARS-CoV-2 infection increases cytoplasmic ERα accumulation and S-ERα colocalization in pulmonary macrophages

To extend the relevance of the above findings to COVID-19, we performed *in vivo* SARS-CoV-2 infection experiments in Syrian Golden hamsters. A SARS-CoV-2/USA-WA1/2020 strain (BEI Resources) was propagated with one passage in cell culture in a biosafety level-3 (BSL-3) laboratory. Syrian Golden hamsters (male, 6-8 weeks old; Envigo, Indianapolis, IN) were exposed to a 1.5 × 10^5^ tissue culture infective dose (TCID50) in 100 μL Dulbecco’s modified Eagle medium (DMEM) by the intranasal route as previously described^31^. Male hamsters were imaged longitudinally inside in-house developed and sealed BSL-3-compliant biocontainment devices^31^ at one day before (Day −1) and at 7 days (Day 7) post-infection using the positron emission tomography (PET) radiopharmaceutical [^18^F]fluoroestradiol ([^18^F]FES) (20 MBq per animal, n = 5) and computed tomography (CT) (**Fig. 4a**). A 90-minute dynamic PET acquisition was performed immediately after intravenous [^18^F]FES injection to visualize the hamster body from the eyes to thighs (starting at the skull vertex). Following PET, a CT scan was immediately performed as previously described^31^. SARS-CoV-2 infection in hamsters produced marked pathology in the lung (as detected by CT and an established image algorithm and analysis pipeline^31^) at Day 7 post-infection compared to Day - 1 (pre-infection) (**Fig. 4b-d**). No distinguishable [^18^F]FES uptake was present in the lungs at Day −1 (**Fig. 4b**). In contrast, the pattern of lung lesions detected via CT overlapped with the lung [^18^F]FES uptake at Day 7 (**Fig. 4b**). Specifically, lung [^18^F]FES uptake at Day 7 was significantly higher in infected lung regions compared to these same sites at Day −1 and at unaffected areas at Day 7 (**Fig. 4c, d**). Furthermore, [^18^F]FES lung uptake at Day 7 was significantly after pretreatment with a pharmacological dose (1 mg/kg, i.v.) of E2, indicating it reflected specific ERα binding (**Fig. 4c, d**). To further corroborate these findings, we performed *ex vivo* biodistribution studies using [^18^F]FES. At 120 min after [^18^F]FES dosing, hamsters were euthanized, and the lungs were harvested and counted for radioactivity. In line with the PET data, SARS-CoV-2-infected hamsters had significantly greater lung [^18^F]FES uptake compared to both uninfected hamsters and SARS-CoV-2-infected hamsters pretreated with E2 (**Fig. 4e**).

**Figure 4.**
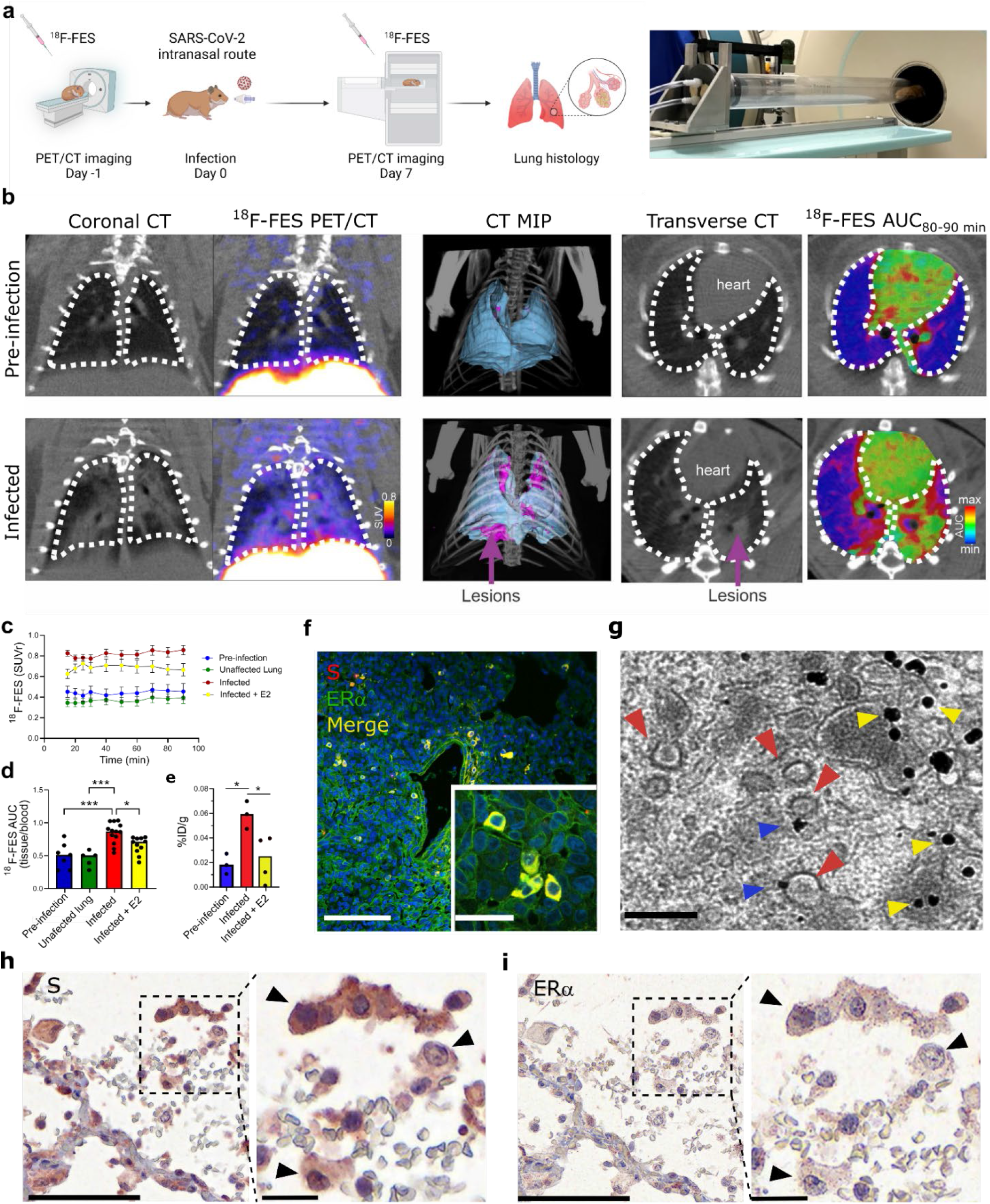
SARS-CoV-2 infection increases cytoplasmic ERα accumulation and S-ERα colocalization in pulmonary macrophages. (**a**) Schematic showing experimental design of SARS-CoV-2 hamster studies and BSL-3 imaging compartment. (**b**) CT, [^18^F]FES PET and AUC heatmap overlay images from hamsters at pre-infection (Day - 1) and infection (Day 7) (MIP; maximum intensity projection, SUV; standard uptake value; AUC, area under the curve). (**c**) Time activity curves showing SUV ratio (SUVr; tissue/blood [^18^F]FES content) in each experimental group. n = 4-5/group. \*\*\**P*<0.001 versus pre-infection (**d**) [^18^F]FES uptake expressed as area under the curve ratio (AUCr; tissue/ blood [^18^F]FES content). F (3, 32) = 12.15; \*\*\**P*<0.001, \**P*<0.05 (**e**) [^18^F]FES uptake expressed as % injected dose (ID)/g body weight in postmortem hamster lung (harvested 110 min post-injection; n = 3-4 /group). F (2, 7) = 7.161; \**P*<0.05. (**f**) Hamster lung immunohistochemistry showing colocalization of S and ERα immunoreactivity. (**g**) Immunogold EM showing SARS-CoV-2 particles (red arrowheads) and ERα-bound gold nanoparticles (blue arrowheads) in a hamster alveolar macrophage (scale bar = 200 nm). Yellow arrowheads correspond to cytoplasmic ERα accumulation. (**h**) S and (**i**) ERα immunostaining in SARS-CoV-2-infected human lung showing S-ERα colocalization in macrophages (black arrowheads). Scale bar (low mag = 100 nm; high mag = 25 nm). All data shown as mean ± SEM.

We exposed additional cohorts of male hamsters to SARS-CoV-2 as above and then sacrificed them at Day 7 post-infection along with uninfected controls and collected their lungs to perform fluorescent immunohistochemistry (IHC) with anti-S and anti-ERα antibodies. As expected, we observed no S or ERα signal in uninfected hamster tissue (**Extended Data Fig. 9**). In contrast, the vast majority of cells from infected hamsters that were positive for S exhibited ERα immunoreactivity (**Fig. 4f and Extended Data Fig. 9**). In infected hamsters, S-positive cells accounted for 14 ± 5%, while ERα-positive cells accounted for 13 ± 5% of lung cells. Moreover, ERα in these cells showed cytoplasmic accumulation in a pattern as we observed in MCF-7 cells transfected with S DNA (**Fig. 3c**).

To examine the subcellular expression of ERα, we performed immunoelectron microscopy (EM) using gold nanoparticles targeting anti-ERα antibodies in lung tissue from uninfected and SARS-CoV-2-infected hamsters. In agreement with the fluorescent IHC results, we found high levels of gold nanoparticle ERα labeling in various cytoplasmic compartments in lung cells from infected hamsters, whereas uninfected hamsters showed low ERα labeling (**Extended Data Fig. 9**). Interestingly, the vast majority of cells from infected hamsters with ERα labeling constituted alveolar macrophages (**Extended Data Fig. 9**) and in such cells, we specifically observed gold nanoparticle accumulation at the surface of SARS-CoV-2 virions (**Fig. 4g**), confirming that S-ERα interact *in vivo*.

Finally, to extend these findings to humans, we performed S and ERα IHC in postmortem lung tissue derived from four human COVID-19 autopsies. We observed S labeling in only one out of the four cases and in that sample S staining colocalized with granular cytoplasmic labeling of ERα in cells favored to be macrophage lineages (alveolar>interstitial) (**Fig. 4h-i, and Extended Data Fig. 10**). The other three cases did not show ERα staining. Importantly, the granular cytoplasmic ERα cytoplasmic staining pattern in the S-labeled cells markedly differed from the nuclear pattern of ERα normally found in breast cancer tissue (**Extended Data Fig. 10**) and was similar to the pattern observed in MCF-7 cells transfected with S DNA and in alveolar macrophage cells in SARS-CoV-2-infected hamsters, supporting the notion that S and ERα colocalize in the cytoplasm of SARS-CoV-2-infected human alveolar macrophage lung cells.

## Discussion

E2 is the most potent endogenous estrogen and is highly selective for ER. In the absence of E2, ERs exist within target cells in a transcriptionally inactive form. Upon ligand activation, ERs undergo homodimerization and binding to discrete DNA regions present at enhancers of specific target genes. Gene regulation occurs when the ER homodimer builds a transcriptional complex with NRC proteins, that can either activate or inactivate transcriptional activity^32,33^. Our results, taken together with prior observations^34^, suggest that S-ERα interactions are involved in SARS-CoV-2 infection and COVID-19 pathology via modulation of ERα signaling, transcriptional regulation of ACE2, and potentially of other genes with roles in inflammation and immunity. Our collective findings indicate that S exhibits structural and functional properties consistent with a role as an NRC at ERα, and it is plausible that this function may extend to other NRs as well. Furthermore, given its conserved LXD motif, it is possible that such properties may also extend to S proteins from other coronavirus strains.

Alveolar macrophages are abundant in the lungs where they play a central role as a first-line defense against various pathogens^35^ including SARS-CoV-2^36,37^. Estrogens are responsible for the maturation and proper functioning of the female reproductive system but also play important roles in immunity^32,38-41^. In particular, ERα signaling in alveolar macrophages is considered a key component of the immune response to infection^42–44^. We observed ER-dependent biological effects of S in the RAW264.7 macrophage cell line. Whereas ERα is mainly localized to the cell nucleus in MCF-7 cells^45^ we found that ERα showed cytoplasmic localization in MCF-7 cells transfected with S DNA. We observed a similar ectopic localization pattern in lung cells, especially in alveolar macrophages from SARS-CoV-2-infected hamsters and humans. More specifically, we found that cytoplasmic ERα co-localized at the surface of SARS-CoV-2 virions within alveolar macrophages, confirming that direct S-ERα interactions occur in the context of SARS-CoV-2 infection in this cell type. Our results, taken together with prior findings, suggest that S-ERα interactions in alveolar macrophages may play a critical role in SARS-CoV-2 infection and COVID-19 pathology.

One of the most frequently reported COVID-19 epidemiologic findings is sex-related mortality and specifically male-related susceptibility. The evidence to date supports a higher predominance of men in several countries; thus, the male sex has been considered a poor prognostic factor^46^. In line with these reports, male laboratory animals are more susceptible to SARS-CoV and SARS-CoV-2 infection and related pathology as compared to females^31,47,48^. ER signaling contributes to these sex differences^48,45^ and the potential protective effects of estrogens in COVID-19 have been widely debated in the literature^49^ though a recent study showed that E2 treatment did not alleviate lung complications in SARS-CoV-2-infected male hamsters^48^. Notably, sex-based differences have been reported in various chronic inflammatory responses associated with lung disease^50^ and specifically, as a function of ER signaling in activated macrophages^42^. Whereas circulating estrogens play a protective role by regulating both the innate and adaptive immune response to infection^51^ it may be possible that the modulation of ER signaling in SARS-CoV-2-infected lung tissue may stimulate proinflammatory signals leading to hypertrophy, vasoconstriction, and vessel obstruction. Indeed, as compared to female patients, hyperactivation of ER signaling in pulmonary tissue in males has been associated with lower frequency but more severe progression of vascular obliteration in pulmonary arterial hypertension^50^. In this context, our data support the notion that S-ERα interactions may lead to an overall dysregulation of ERα signaling and lung lesion development. Our results account for a model in which S-mediated tissue-specific dysregulation of ERα signaling is a key event in COVID-19 lung pathogenesis that may contribute to SARS-CoV-2 development in specific categories of subjects/patients with low basal ER signaling such as men and post-menopausal women. This model could also potentially explain the widely discussed effect of ER modulation in SARS-CoV-2 infection and the reported protective effect of anti-estrogenic treatment on COVID-19 prevalence in women with ovarian and breast cancer^4^

In conclusion, we report novel interactions between the SARS-CoV-2 S and ERα that may have important therapeutic implications for COVID-19. Our results also highlight the use of multimodal PET/CT imaging and the FDA-approved [^18^F]FES radiopharmaceutical as a translational approach and biomarker for the longitudinal assessment of COVID-19 lung pathology. Finally, we propose that tissue-specific dysregulation of ER activity should be considered in the design of S-based vaccines.

## Acknowledgements

This work was supported by the National Institute on Drug Abuse (ZIA000069) to MM, the R01-AI153349, R01-AI161829-A1, and the Center for Infection and Inflammation, Johns Hopkins University to SJ, and the National Institute of Biomedical Imaging and Bioengineering P41 (EB024495) to MGP. We thank Nicolai Urban (Max Planck Institute for Neuroscience Imaging Center), Mario Borgnia (National Institute on Environmental Health Sciences), Lei Shi (National Institute on Drug Abuse), Silvano Coletti (Chelonia Applied Science), Matthew Hall (National Center for Advancing Translational Sciences) and Alessandro Grottesi (Department HPC, CINECA, via dei Tizii 6, 00185 Roma, Italy) for sharing reagents, technical advice, and discussions. MM has received research funding from AstraZeneca, Redpin Therapeutics, and Attune Neurosciences. All other coauthors report no conflicts of interest.

## Material and Methods

### SARS-CoV-2 S radiolabeling

To an eppendorf vial was added 0.25 M phosphate buffer pH 7.5 (80 μL), SARS-CoV-2 S (R683A, R685A), His Tag (20 μg) (Acro Biosystems, #SPN-C52H4) in water, lactoperoxidase (2 μg), Na125I (0.7 mCi), H_2_O_2_ (0.4E-03%). After incubation for 60 minutes at 35°C the reaction was quenched with ascorbic acid (0.1 mg). The mixture was allowed to stand for 10 minutes, and then bovine serum albumin (3 mg) was added. The mixture was then applied to a G-25 desalting column (GE Healthcare) to separate the radioiodinated S from unreacted radioiodine. Approximately 0.2 mCi of product was obtained. The purified radiolabeled protein was formulated with 1% BSA and 10% sucrose, divided into aliquots and stored at −20 °C.

### [^125^I]S radioligand binding assays

For saturation assays, the radioligand specific activity was adjusted with unlabeled peptide to enable a radioligand concentration range appropriate for the Kd of the receptor. Plate preparation: Protein A-coated plates (ThermoScientific Cat. # 15130) were washed with wash buffer (50 mM Tris, 5 mM MgCl2, 0.1 mM EDTA, pH 7.4) and incubated with human ACE2 Fc Tag (0.2 μg/well) (Acro Biosystems, #AC2-H5257) in incubation buffer (50 mM Tris, 5 mM MgCl2, 0.1 mM EDTA, 2% BSA, pH 7.4) for 60 minutes with gentle shaking. The plates were then washed with wash buffer (4 x 0.3 ml) followed by high salt wash buffer (50 mM Tris, 5 mM MgCl2, 0.1 mM EDTA, 125 mM NaCl, pH 7.4) and used directly for the binding assay. Incubation and filtration: To each well was added 75 μL buffer, 25 μL of the unlabeled protein (for competition assays) or buffer and 25 μL of radioligand solution in binding buffer. The plate was incubated at RT for 60 minutes with gentle agitation. The incubation was stopped by washing the wells with (i) incubation buffer (1 x 0.3 ml, ice cold), (ii) wash buffer (3 washes, ice cold) and (iii) high salt wash buffer (1 wash; ice cold). Following washing, to each well was added NaOH (0.1 M) and the plates were incubated at 40°C for 1 hour to digest the protein. Following digestion, the radioactivity was transferred to a counting plate, neutralized and scintillation cocktail (Betaplate Scint; PerkinElmer) added and the radioactivity counted in a Wallac^®^ TriLux 1450 MicroBeta counter. For each concentration of radioligand, non-specific binding was subtracted from total binding to give specific binding. Non-specific binding was determined using wells incubated without ACE2. For saturation assays, the bound radioactivity (CPM/well) was converted to molar amounts (fmol/well) from the specific activity of the radioligand. A counting efficiency for 125I of 71% was used for CPM. to DPM. calculations. Data were fitted using the non-linear curve fitting routines in Prism^®^ (Graphpad Software Inc).

### Protoarray

The Invitrogen ProtoArray^®^ Human Protein Arrays (ThermoFisher Scientific) are high-density microarrays that contain more than 9,000 unique human proteins individually purified and arrayed onto a nitrocellulose-coated slide. We followed the manufactured instructions to probe the arrays for small tritiated molecules. Briefly, protein microarrays were blocked for 30-40 min in blocking buffer (50 HEPES, 250 NaCl, 20 glutathione, 1 DTT, 1% (or 2%) BSA, 0.1% Tween). The blocking buffer was then gently aspirated off and replaced with incubation buffer (Phosphate buffered saline, 0.1% Tween, 1% (or 2 %) BSA, w/wo 1 nM 3H-E2) containing the radioligand ([125I] S (20 nM)). To determine non-specific binding, 300 nM of S was added to the incubation mix. Every condition was tested in duplicate. After incubation, slides were washed 3 times in ice-cold washing buffer (Phosphate buffered saline, 0.1% Tween) and rinsed with ice-cold distilled water. Slides were then air dried and placed into a Hypercassette™ and covered by a tritium-sensitive phosphor screen (GE healthcare), exposed for 1 day and then scanned on a PerkinElmer Cyclone^®^ scanner. The digitized images were also analyzed using ProtoArray Prospector v5.2 and potential hits were identified using the software’s algorithm.

### Surface plasmon resonance using immobilized SARS-CoV-2 S

SPR measurements were performed using a Biacore apparatus (Biacore) using CM5 sensor chips. To find out the optimal pH for S (Acro Biosystems) immobilization, we conducted pH scouting. The S was prepared in 10 mM sodium acetate buffer at pH 4.0 to 5.5. The best pH for immobilization was 4.0 (**Extended Data Fig. 11a**). After covalent immobilization there was approximately 8500 RUs of S on the sensor surface (**Extended Data Fig. 11b**). Increasing concentrations of ERα-full length (InVitrogen), ACE2 (Acro Biosystems) and NRP1 (Acro Biosystems) from 1.56 to 200 nM were injected. Protein binding responses were analyzed using BiaEval software. all curves were globally fitted to a single site binding model to determine an approximate fit. The Chi2 value was noted to indicate the goodness of fit. The remaining data were refitted to the single site binding model and the improvement in fit (reduction in Chi2 value) noted. K_on_, K_off_ and KD were reported by the global fit model. These data were also fitted to a 2-site binding model to determine if this gave a better fit (indicated by a lower Chi2 value than the single site model). For the equilibrium model, binding response amplitudes were fitted to a saturation binding curve, from which the Rmax and KD (concentration at half Rmax) were determined.

### Interactome analysis

The STRING database^52^, that integrates all known and predicted associations between proteins, including both physical interactions as well as functional associations has been used to analyses functional associations between biomolecules. Each protein-protein interaction is annotated with a ‘scores’. This score does not indicate the strength or the specificity of the interaction but the confidence. All scores rank from 0 to 1, with 1 being the highest possible confidence.

### 3D Model Selection and MD simulation protocol

S 3D model was built based on PDB 6VYB returned to its wild-type form and fully glycosylated^53^. An asymmetric glycosylation of the three protomers has been derived by glycoanalyitic data for the N-glycans and O-glycans according to the work of ^54,55^. For the estrogen receptor, the X-RAY PDB model with code 3UUD was used, containing E2α and Nuclear receptor coactivator 2^23^ and 3OLL, containing E2β and Nuclear receptor coactivator 1^24^. The proteins were modeled using Amber14SB force field^56^ and the carbohydrate moieties by the GLYCAM06j-1 version of GLYCAM06 force field^57^ and the general amber force field (GAFF)^58^ was used for the estradiol bound to ER receptor. The so prepared structure was used as starting point for MD simulations. Protein was inserted in a clinic box, extending up to 10 Å from the solute, and immersed in TIP3P water molecules^59^. Counter ions were added to neutralize the overall charge with the genion GROMACS tool. After energy minimizations, the system was relaxed for 5 ns by applying positional restraints of 1000 kJ mol-1 nm^-2^ to the protein atoms. Following this step, unrestrained MD simulation was carried out with a time step of 2 fs, using GROMACS 2020.2 simulation package (supercomputer Marconi-100, CINECA, Bologna, Italy)^60^. V-rescale temperature coupling was employed to keep the temperature constant at 300 K^61^. The Particle-Mesh Ewald method was used for the treatment of the long-range electrostatic interactions^62^. The first 5 ns of each trajectory were excluded from the analysis. The trajectory obtained after 1 microsecond MD simulation has been clustered in order to obtain representative structures. In particular, the structure used for the docking studies is the first centroid of the first cluster extracted from the MD experiment.

For the ER, the XRAY PDB model with code 3OLL was used, containing E2 and Nuclear receptor coactivator 1^24^.

### Protein-Protein Docking procedure

The input of two individual proteins, one for receptor and the other for ligand, were provided. In particular, the S and ER was used as receptor and ligand respectively. Then, the HDOCK tool will perform docking to sample putative binding modes through an FFT-based search method and then scoring the protein–protein interactions. Finally, the top 100 predicted complex structures are provided, and the best ten hypotheses were visually inspected to confirm the reliability of the calculation. The entire workflow is well described in the work published by Yan et al.^26^

### ERα reporter gene assays

#### Peptide transfection

GeneBLAzer^®^ ERa-UAS-bla GripTite™ (HEK293 ERa-bla; Invitrogen, Carlsbad, CA, USA) cells comprise a mammalian one-hybrid system stably expressing a beta-lactamase reporter gene under the control of the GAL4 DNA-binding site and a fusion protein consisting of the human ERα ligand-binding domain and the GAL4 DNA-binding domain. The cells were plated at 8,000 cells/ well in black wall/ clear bottom plate. After cultured overnight, cells were treated with peptides delivery reagent, 20mM HEPES containing peptides and PLUSin^®^ reagent (Polyplus-transfection, NY USA), for 4 hours in DMEM supplemented with 1μM NEAA and 100U/mL penicillin, 100μg/mL streptomycin. Then, medium was replaced with phenol-red free DMEM supplemented with 2% charcoal stripped FBS, 1μM NEAA and 100U/mL penicillin, 100μg/mL streptomycin, and 10μM sodium pyruvate.

#### S transfection

One million cells were plated into each well of 6-well plates in 2 ml of DMEM supplemented with 10% FBS, 1μM NEAA and 100U/mL penicillin, 100μg/mL streptomycin. On the next day, the medium was replaced with 2 ml of DMEM supplemented with 10% FBS. Then, the cells in each well were transfected with 2.5 μg of different spike plasmids with Lipofectamine 3000. After 24 hours of incubation, cells were detached from 6 well plate using tryptophan and plated at 8,000 cells/ well in black wall/ clear bottom 384 well plate.

Transfected cells were treated with E2 for 18 hours. The next day, 8 μL of LiveBLAzer™ (Life Technologies, Madison, WI) detection mixture was added to each well and the plates were incubated at room temperature in the dark for 2 h. Fluorescence intensity at 460 and 530 nm emission and 405 nm excitation was measured by an PHERAstar plate reader (BMG LABTECH, Cary, NC). Data were represented as the ratio of the emission wavelengths (460nm/530nm).

### Human ERα transcriptional activation assays

The ERα activity was determined by the ERα transcription factor activation assay kit (ab207203, Abcam) according to manufacturer’s directions. Briefly, MCF-7 nuclear extracts (5 μg; ab14860, Abcam) were treated with either S (0.01-300 nM; Acro Biosystems) S-RBD (1-100 nM; Acro Biosystems), S-trimer (1-100 nM; provided by Dr. Borgnia) and/or E2 (100 nM: Tocris). Extracts were added to each well coated with the ER consensus binding site (5’ – GGTCACAGTGACC – 3’). The wells were washed and then incubated with rabbit anti-ERα (1:2000, 1 h, RT) and horseradish-conjugated secondary antibody (1:2000, 1 h, RT) that were provided with the kit. Colorimetric reaction was measured by spectrophotometry at a wavelength of 450 nm.

### Immunofluorescence (IF)

About 80,000 MCF-7 cells were placed in each chamber of a 4-well chamber slide (Thermo Scientific, cat. no. 177399) containing 500 μl of Dulbecco’s Modified Eagle Medium (DMEM) + 10% fetal bovine serum (FBS) and cultured overnight at 37°C in a 5% CO2 incubator. The next day, cells in each well were transfected with 1.5 μl of ViaFect reagent (Promega, cat no. E498A) and 0.5 μg of empty pcDNA3.1 vector, or an expression vector for the wild-type (WT) SARS-CoV2 S with a C-terminal hemagglutinin (HA) epitope tag (pBOB-CAG-SARS-CoV2-S-HA) or the double mutant (R682S,R685S) SARS-CoV2 S with a C-terminal flag epitope tag (pCAGGS-SARS2-S-FKO). pBOB-CAG-SARS-CoV2-S-HA was a gift from Gerald Pao (Addgene plasmid # 141347; http://n2t.net/addgene:141347; RRID:Addgene_141347). pCAGGS-SARS2-S-FKO (C-flag) was a gift from Hyeryun Choe & Michael Farzan (Addgene plasmid # 159364; http://n2t.net/addgene:159364; RRID:Addgene_159364). After 48 hours, the cells were fixed in 4% formaldehyde for 20 min at room temperature, rinsed with 1X phosphate-buffered saline (PBS), permeabilized with 0.1% Triton X-100 for 25 min, rinsed with PBS, and incubated for 2 hours in blocking buffer (Rockland Immunochemicals, Inc. cat no. MB-070). The cells were then incubated at 4°C overnight with 2 μg/ml each of anti-ERα(H222) rat IgG1 monoclonal antibody (mAb) (Santa Cruz Biotech, sc-5349, 1:100) and HA-probe (F-7) mouse IgG2a mAb (Santa Cruz Biotech, sc-7392X, 1:1,000), OctA-probe (anti-flag) mouse IgG1 mAb (Santa Cruz Biotech, sc-51590, 1:50), or normal mouse IgG (Santa Cruz Biotech, cat no. sc-2025, 1:200) as a negative control. Afterwards, the cells were washed 4 times with PBS + 0.1% Tween-20 (PBS-T) for 5 minutes and incubated at room temperature for 1 hour in the dark with a fluorescent secondary antibody mixture contaning mouse IgGk BP-CFL594 (Santa Cruz Biotech, sc-516178, 1:100) and anti-rat IgG AF488 (ThermoFisher Scientific, cat no. A-11006, 1:500). The cells were then washed 4 times with PBS-T for 5 minutes in the dark and rinsed with PBS. Each slide was carefully detached from its gasket, and immediately mounted with a 1.5T glass coverslip using EverBrite Hardset Mounting Medium with DAPI (Biotium, cat no. 23004). The mounted slides were allowed to cure for 24 hours in the dark at room temperature, and stored in a slide box at 4°C. The slides were imaged at 63X magnification using a Zeiss LSM 880 Airyscan inverted confocal microscope (Max Planck Florida Institute for Neuroscience). Each image represents the average of 16 scans. Images were prepared for presentation using ImageJ v.1.53c software (NIH).

### Proliferation assays

MCF-7 and MDA-MB-23 cells were obtained from ATCC and growth in DMEM without phenol red, supplemented with 10% fetal bovine serum (FBS), penicillin/streptomycin at 37 °C in a 5% CO2 and 95% humidified atmosphere. For each assay cells were seeded at the density of 10^4^ cells/cm^2^. Before treatments, to reduce estrogen levels in FBS and avoiding any interference, cells were cultured for 24h in medium containing 5% dextran-coated charcoal treated serum. Then, cells were treated for 24h with E2 (Sigma-Aldrich; Cat: E1024; Batch: SLCC8875), S (R&D Systems; Cat: 1059-CV; Batch: DODR0220111), raloxifene (Sigma-Aldrich; Cat: R1402; Batch: MKCJ7180), S + raloxifene, E2 + S, E2 + raloxifene + S. In particular, the concentration tested for S was 10 ng/ml, for raloxifene 2 μM, for E2 1 nM.

Cell proliferation was measured using a 5-bromo-2-deoxyuridine (BrdU) labeling and a proliferation ELISA Kit (Abcam, ab126556) following the manufacturer’s instructions. Briefly, BrdU was added to wells for 24h and then cells were fixed using Fixing Solution. Then, cells were washed and were incubated with detector anti-BrdU antibody for 1 hour at RT. After the incubation cells were washed and incubated with the horseradish peroxidase conjugated goat anti-mouse antibody for 30 minutes at RT. For the detection the chromogenic substrate tetra-methylbenzidine (TMB) was added and the colored product has been detected using a spectrophotometer (450/550 nm). Values were given as percentage of cells grown only in serum-free medium. At least two independent assays were performed with eight duplicates each.

### TRAP activity by ELISA assay in RAW-OCs

RAW264.7 (murine macrophages ATCC, USA) were cultured as manufacturer’s protocol. Then 1.5x 10^5^ cells/cm^2^ in 24-well dishes were seeded and mouse receptor activator of nuclear factor kb ligand (RANKL, Miltenyi Biotec, Germany) was added at the final concentration of 35 ng/ml to initiate osteoclasts (OC) development (day 0) as previously described^63^. At day 3, cells were examined under the microscope and refed with fresh medium containing RANKL. At day 6, RAW-OC population was prevalent and ready for treatments and then biochemical studies. Cells were treated with E2 (1 nM), S (10 ng/ml) and raloxifene (2 μM) and the combination of them for 24h. After 24 h of treatment, we quickly collect the cells by sterile tubes and resuspended the cells using PBS (pH 7.4) to dilute cell suspension to the concentration of approximately 1 million/ml. Then, cells were subjected to repeated freeze-thaw cycles to let out the inside components. In the meantime, the reagents of the kit were brought to room temperature.

Tartrate Resistant Acid Phosphatase (TRAP) activity was performed using an ELISA kit from Mybiosource (MBS1601167). The standard curve, reagents and samples were prepared following manufacturer’s protocol. Briefly, 50 μl of standard were added to standard wells and 40 μl of sample-to-sample wells and then added 10 μl of anti-TRAP antibody to sample wells and 50 μl of streptavidin-HRP to sample wells and standard wells. The plate was incubated 1 hour at 37°C. The plate was washed 5 times with wash buffer and 50 μl of substrate solution A were added to each well plus 50 μl of substrate solution B and incubated 10 minutes at 37°C in the dark. Finally, 50 μl of stop solution to each well were added and the optical density was immediately determined using a microplate reader set at 450 nm.

### Fluorescence-based assay for ACE2 in MCF-7 cells

Cells were obtained from ATCC and grew in DMEM without phenol red, supplemented with 10% fetal bovine serum (FBS), penicillin/streptomycin at 37 °C in a 5% CO2 and 95% humidified atmosphere. For each assay cells were seeded at the density of 10^4^ cells/cm^2^. Before treatments, to reduce estrogen levels in FBS and avoiding any interference, cells were cultured for 24h in medium containing 5% dextran-coated charcoal treated serum. Then, cells were treated for 24h with E2, S, raloxifene, S + raloxifene, E2 + S, E2 + raloxifene + S. In particular, the concentration tested for S was 10 ng/ml, for raloxifene 2 μM, for E2 1 nM.

Cells were cultured in a 96-well plate. Cells were washed in ice-cold PBS and then fixed with 2% Formalin solution in PBS for 15 minutes. After further washes, to prevent the non-specific binding, cells were blocked with a 10% Bovine Serum Albumin solution in PBS for 20 minutes and then incubated with the Alexafluor 647-conjugated antibody for Human ACE-2 (R&D Systems). After several washes, the plate was read at the fluorescence intensity of 668 nm using a microplate reader (Spark, Tecan). Then, to normalize the results DAPI was added and the fluorescence intensity at 461 nm was evaluated.

### ACE2 expression in Calu-3 cells

Calu-3 cell line was obtained from ATCC and maintained in Eagle’s Minimum Essential *Medium*(EMEM; Lonza) supplemented with 10% fetal bovine serum (FBS), 1% L-glutamine and 1% penicillin/streptomycin solution at 37°C in a humidified atmosphere of 5% CO2.

#### RNA extraction and qRT-PCR

Cells were seeded in 6 well plates and after incubation, were treated according to the experimental protocol with E2 (200nM), Raloxifene (20μM), S (10ng/ml). Total RNA was extracted from cell lines using RNeasy^®^ Plus Mini Kit (Qiagen). cDNA was made using GenePro thermal cycler (Bioer). RT-PCR analysis was performed on an Applied Biosystem™ QuantStudio™ 5 Real-Time PCR system (Thermo Fisher Scientific) using Itaq™ Universal SYBR (Bio-Rad) gene expressions assays. Primers for GAPDH (forward primer AATCCCATCACCATCTTCCA; reverse primer TGGACTCCACGACGTACTCA) and ACE2 (forward primer AAAGTGGTGGGAGATGAAGC; reverse primer GAGATGCGCGGTCACAGTAT) were used. Samples were assayed in runs which were composed of 3 stages: hold stage at 95°C for 20 minutes, PCR stage at 60°C for 25 minutes and melt curve stage 95°C for 1 minute, 60°C for 20 minutes, and 95°C for 1 minute again. Gene expressions were normalized by GAPDH levels using the 2-ΔΔCt method.

#### Immunocytochemistry assay

Cells were seeded on glass coverslips pre-coated with collagen in 24-well plates. After incubation at 37°C, cells were treated according to the experimental protocol with E2 (200nM), Raloxifene (20μM), S (10ng/ml). After 72 hours, cells were washed, fixed with 4% formaldehyde, permeabilized with 0.1% Triton X-100 in PBS and stained overnight at 4°C with ACE2 protein-specific antibody (Abcam Ab15348). Cells were then incubated with anti-rabbit secondary antibody (Alexa Fluor 536 anti-rabbit, Invitrogen Life Technologies) for 1 hour at 37°C. Nuclei were labeled with Hoechst 33342 (Thermo Fisher Scientific) for nuclear staining for 20 minutes.

Cells were mounted with Fluor-mount (Sigma-Aldrich, St Louis, MO, USA) and images were acquired through confocal microscope LSM 800, magnification 60X, software ZN 2.1 blue Edition (Carl Zeiss, Jenza, Germany) and analyzed with ImageJ software.

### Positron emission tomography (PET) imaging

#### Animal studies

All animal protocols were approved by the Johns Hopkins University Biosafety, Radiation Safety, and Animal Care and Use Committees. Male golden Syrian hamsters (7 to 8 weeks of age) were purchased from Envigo (Haslett, MI). Animals were housed under standard housing conditions in positive/negative control cages (PNC) (Alentown, NJ) in animal biological safety level 3 (ABSL-3) facility at the Johns Hopkins University-Koch Cancer Research Building. After 1-2 weeks of acclimation, animals were inoculated with 1.5 x 10^5^ TCID50 of SARS-CoV-2 USA-WA1/2020 in 100 μL of DMEM (50 μL/nostril) through the intranasal route under ketamine (60 to 80 mg/kg) and xylazine (4 to 5 mg/kg) anesthesia administered intraperitoneally, as previously described^43^. Control animals received an equivalent volume of DMEM.

#### Imaging

Hamsters were imaged inside in-house developed; sealed biocontainment devices compliant with ABSL-3. A cohort of male non-castrated hamsters were imaged longitudinally one day before SARS-CoV-2 infection and 7 days post-infection using [^18^F]fluoroestradiol (FES) (n = 4-5). A second cohort of SARS-CoV-2-infected male hamsters were intravenously co-injected with 18F-FES and 0.3 mg/kg of E2 (3% DMSO; Sigma-Aldrich) on day 7 post-infection (n = 4). Each animal was injected 16.1 ± 1.5 MBq of [^18^F]-FES intravenously via the penile vein. A 90-minute PET acquisition and subsequent CT were performed using the nanoScan PET/CT (Mediso, Arlington, VA). For each animal, eight to thirteen volumes of interest (VOIs) were manually selected using CT as a guide and applied to the PET dataset using VivoQuantTM 2020 (Invicro, Boston, MA) for visualization and quantification. A VOI was placed on the left ventricle of the heart to measure the blood uptake. [^18^F]-FES PET activity was calculated for each hamster (n = 4-5 hamsters per group) as the average activity of all VOIs normalized by the mean standardized uptake value (SUVmean). All animals were sacrificed 110 min post-injection and the lung harvested to quantify associated radioactivity using an automated γ-counter. Heatmap overlays were implemented using RStudio Version 1.2.1335 (R Foundation) as previously described^64^. Multiple comparisons were performed using two-way repeated measures analysis of variance (ANOVA) followed by Bonferroni’s multiple-comparison test.

### Hamster histology

Hamster Lung Sample Preparation for Confocal and Electron Microscopy: We perfused uninfected and infected hamsters with 1000U/ml heparin solution followed by fixative solution (4% paraformaldehyde (PFA), 0.15% glutaraldehyde, 15% picric acid solution in 0.1 M phosphate buffer, pH=7.4 (PB)) or fixative solution (4% paraformaldehyde (PFA), 15% picric acid solution in 0.1 M phosphate buffer, pH=7.4 PB). After perfusion, kept the lungs in the same fixative solution at 4°C for another 2 h. We placed the lungs in 2% PFA fixative solution and post-fixed the lungs at 4°C overnight. After rinsing in 0.1 M PB, serial sections (50 μm) were cut with a vibratome (VT1000S, Leica Microsystems Inc).

#### Immunohistochemistry staining in hamster tissue

The vibratome lung sections were rinsed and incubated for 1 h in 0.1 M PB supplemented with 4% BSA and 0.3% Triton X-100. Sections were then incubated with cocktails of primary antibodies: rabbit anti-SARS-CoV-2 Spike Protein (1:100, Invitrogen, #MA5-36087) + rat anti-ERα H222 (1:100, Santa Cruz Biotechnology, #sc53492) overnight at 4°C. After rinsing 3 × 10 min in PB, sections were incubated in a cocktail of the corresponding fluorescence secondary antibodies: Alexa-Fluor-594-donkey anti-rabbit (711-585-152, Jackson ImmunoResearch Laboratories) + Alexa-Fluor-488-donkey anti-rat (712-545-153, Jackson ImmunoResearch Laboratories) for 2 h at room temperature. After rinsing, sections were mounted on slides. Fluorescent images were collected with a Zeiss LSM880 with Cy7.5 Confocal System (Zeiss). Images were taken sequentially with different lasers with 20× objectives.

#### Electron Microscopy

The vibratome lung sections were rinsed and incubated with 1% sodium borohydride to inactivate free aldehyde groups, rinsed, and then incubated with blocking solution. Sections were then incubated with the primary antibodies rat anti-ERα H222(1:100, Santa Cruz Biotechnology, #sc53492), diluted in 1% normal goat serum (NGS), 4% BSA, 0.02% saponin in PB at 4°C overnight. Sections were rinsed and incubated overnight at 4°C in the secondary antibody Nanogold-Fab’ goat anti-rat-IgG (1:100, Nanoprobes, #2008) for ERα protein detection. Sections were rinsed in PB, and then sections were post-fixed with 1.5% glutaraldehyde for 10 min and rinsed in PB and double-distilled water, followed by silver enhancement of the gold particles with the Nanoprobe Silver Kit (2012, Nanoprobes) for 7 min at room temperature. Sections were rinsed with PB and fixed with 0.5% osmium tetroxide in PB for 25 min, washed in PB, followed by ddH2O water, and then contrasted in freshly prepared 1% uranyl acetate for 35 min. Sections were dehydrated through a series of graded alcohols and with propylene oxide. Afterwards, they were flat embedded in Durcupan ACM epoxy resin (14040, Electron Microscopy Sciences). Resin-embedded sections were polymerized at 60°C for 2 days. Sections of 60 nm were cut from the outer surface of the tissue with an ultramicrotome UC7 (Leica Microsystems) using a diamond knife (Diatome). The sections were collected on formvar-coated single slot grids and counterstained with Reynold’s lead citrate. Sections were examined and photographed using a Tecnai G2 12 transmission electron microscope (Thermo Fisher Scientific) equipped with the OneView digital micrograph camera (Gatan).

### Serial Immunohistochemistry Staining in human tissue

Formalin fixed and Paraffin embedded human breast and human covid lung tissues were achieved from Pathology Department of Johns Hopkins Medicine under an IRB-approved protocol. The tissue slides were deparaffinized with xylene and rehydrated with gradient concentrations of ethanol; boiled in a high-pressure cooker with a citrate buffer (BioSB Inc., Catalog No. BSB 0032) for 15 minutes retrieval. Then, slides were subjected to the serial immunohistochemistry. On day one, slides were blocked with a peroxidase blocker (Bio SB Catalog No. BSB 0054), washed with an immunoDNA washer buffer (Bio SB, Catalog No. BSB 0150); then, incubated with 0.2 μg/mL of anti-SARS-CoV-2 spike glycoprotein antibody (abcam, Catalog No. ab272504) for 1 hour. After three washes, the Mouse/Rabbit PolyDetector Plus link &HRP label (Bio SB, Catalog No. BSB 0270) were applied. The AEC-red chromogen (Vector Laboratories, catalog No. SK-4205) and a Hematoxylin solution were used for color development and countered stain. The slides were mounted with a VectaMount AQ Aqueous Mounting medium (Vector Laboratories, Catalog No. H-5501) and scan at 20x and 40x magnification using a MoticEasyScan Pro 6 (Meyer instruments Inc. Houston TX). After scan, the slides were incubated in a PBS buffer for cover slip detachment. Day two, the slides of detached coverslip were dehydrated in 70% Ethanol; decolorized in 90% Ethanol for 10 min; rehydrated with 70% Ethanol and dH2O. The decolorized slides were stripped with an antibody elution buffer (0.2% SDS, 62.5mM Tris-HCI, pH 6.8, 5% Glycerol and 0.08%β-mercaptoethanol) within a 130 °C oven for 20 min (until boiling). The slides were washed with dH2O three times, each 15 minutes; wash buffer twice, each 10 minutes; and then incubated in PBS buffer for 10 minutes. After antibody elution, an Estrogen-α antibody (abcam, Catalog No. ab108398) at 1:250 work solution was applied for 90 min, and then followed previous procedure for stain development. Finally, the slides were mounted and scanned for data collection. Images were quantified with the “cell counter” plug-in in ImageJ (NIH).

