## Extended Data for "The SARS-CoV-2 spike protein binds and modulates estrogen receptors"

**Extended Data Figure 1**

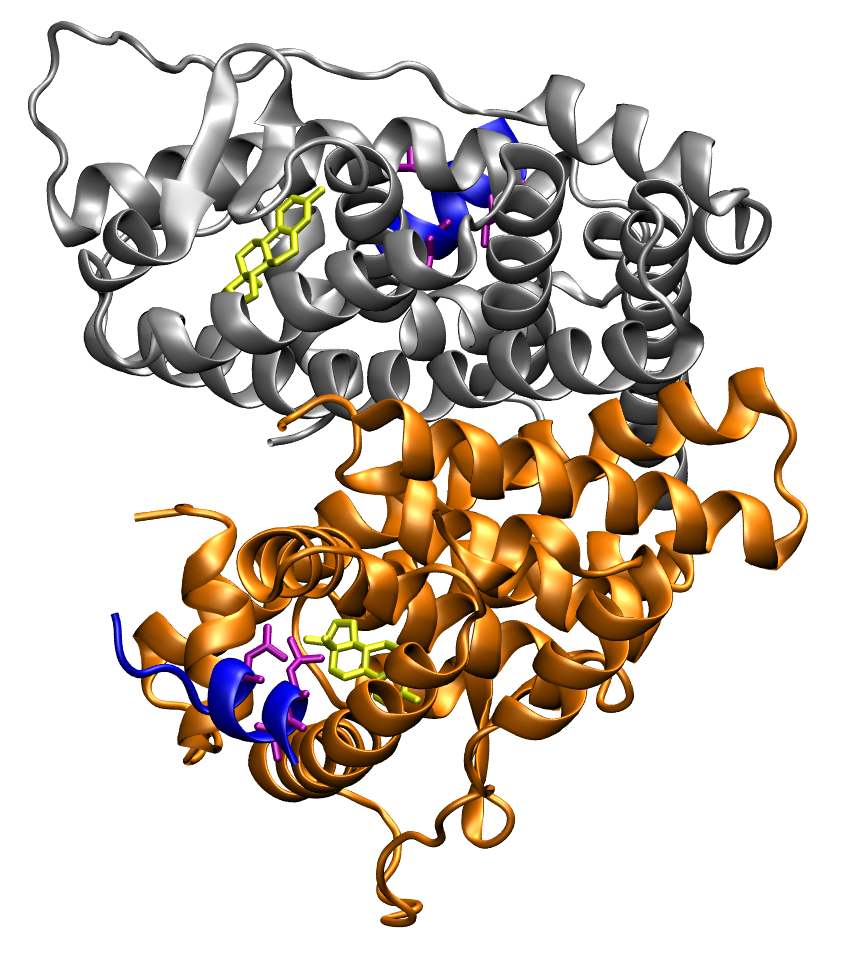

**Extended Data Figure 1.** Crystal structure of ERα dimer in interaction with a NCOA1 fragment containing the LXD nuclear receptor coregulator (NRC) motif (pdb id 3UUD). The AF2 region in ERα (residues 302-552) is shown in silver and orange colors for the two monomers. The NCOA1 fragment containing the LHKLL (residues 690-694) is represented in blue color. Lateral chains of the three leucine residues are also represented in purple color. Estradiol is shown in yellow color.

**Extended Data Figure 2.**

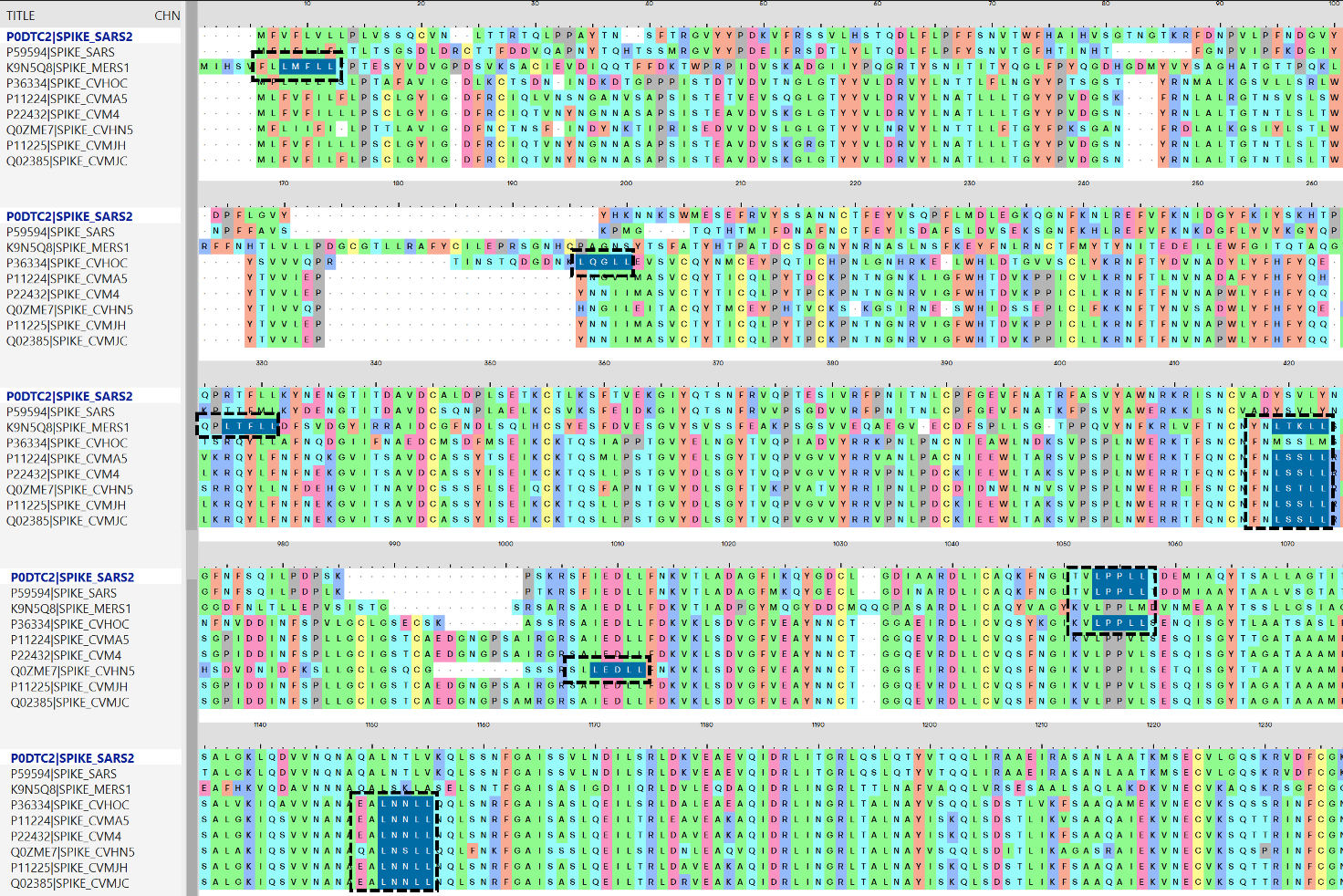

**Extended Data Figure 2.** S protein sequence alignment and LDX motif identification. Amino acid patterns related to the LDX-like motif, conserved among different species, are highlighted in blue, and surrounded by black dotted boxes.

**Extended Data Figure 3.**

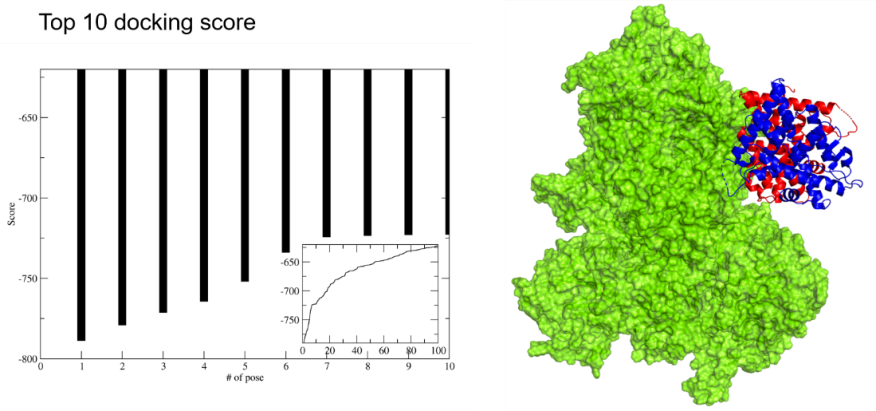

**Extended Data Figure 3.** Binding hypothesis of ER to S by guided docking study. On the left side the plot reporting the top 10 docking scores and the score distribution along the 100 structures generated by the docking software (the smaller plot below). On the right side, the best 3D docking hypothesis. The ER dimer is in blue and red cartoon, S surface is shown in green.

**Extended Data Figure 4**

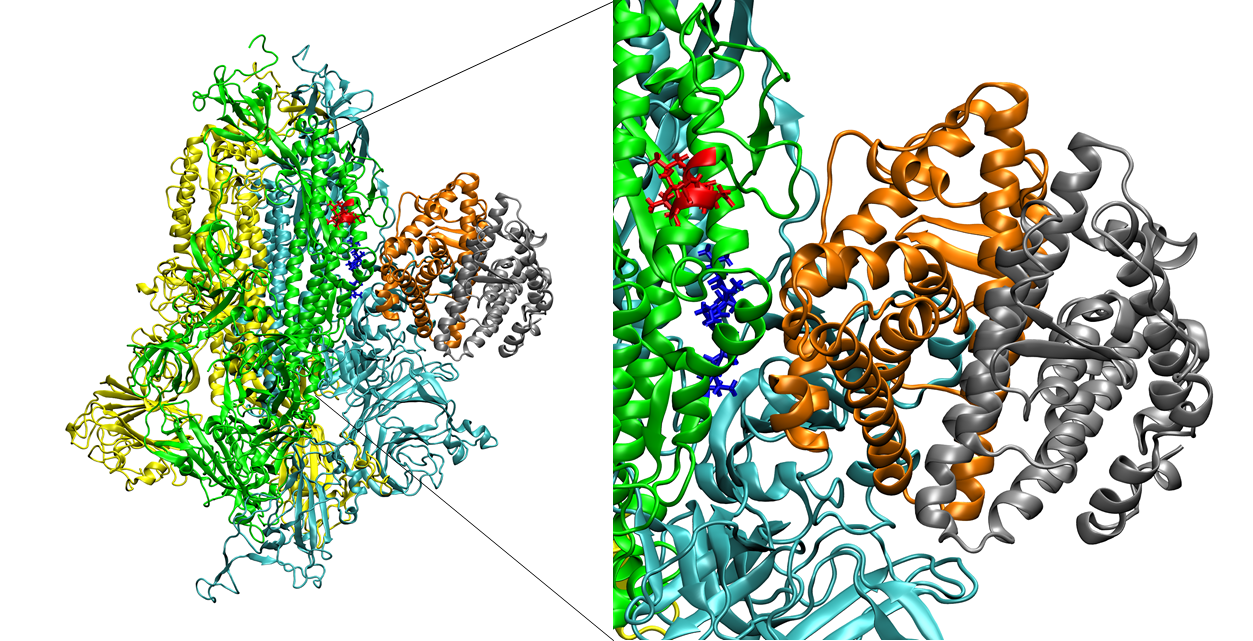

**Extended Data Figure 4.** Snapshot of molecular dynamics frame for the S-ERα complex. The simulation shows the formation of a strong interaction surface between ER and S, in the region of the two conserved LXD motifs. The average number of protein-protein direct hydrogen bonds per timeframe is more than 14 after only few hundred nanoseconds.

**Extended Data Figure 5.**

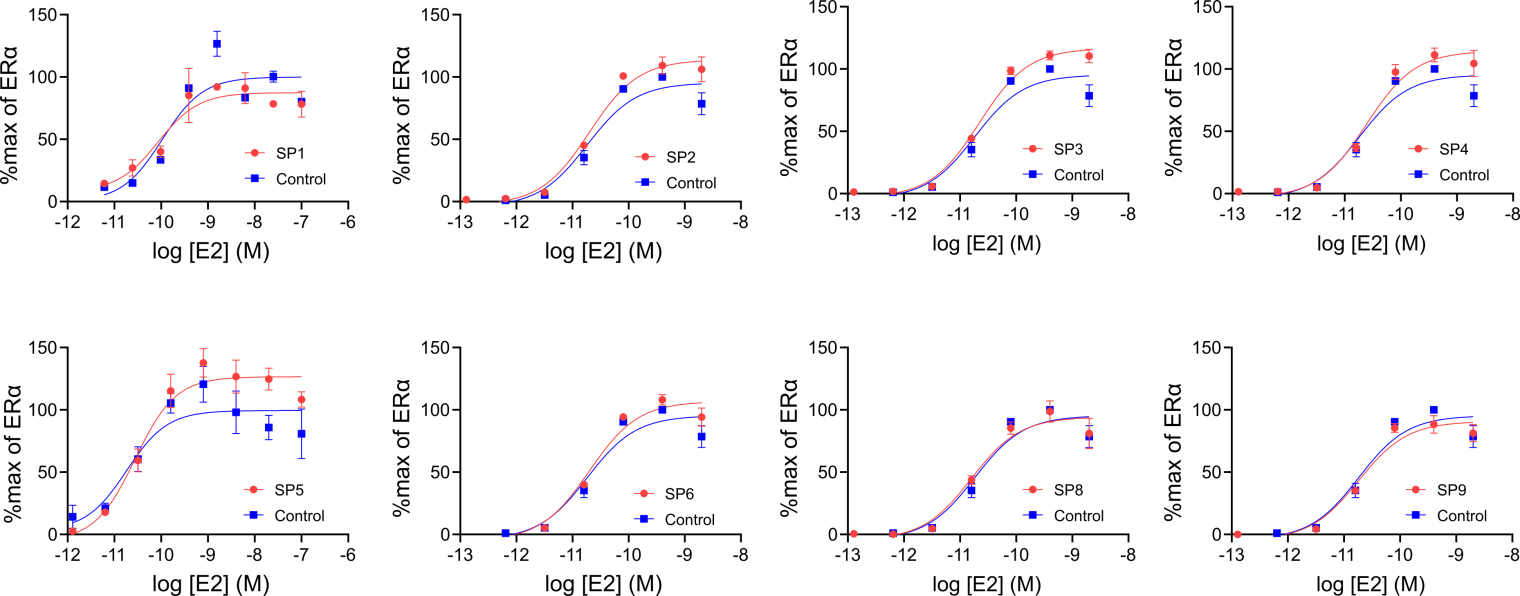

**Extended Data Figure 5.** Effect of S peptides on ERα-mediated transcriptional activation. Data are shown as mean ± SEM.

**Extended Data Figure 6.**

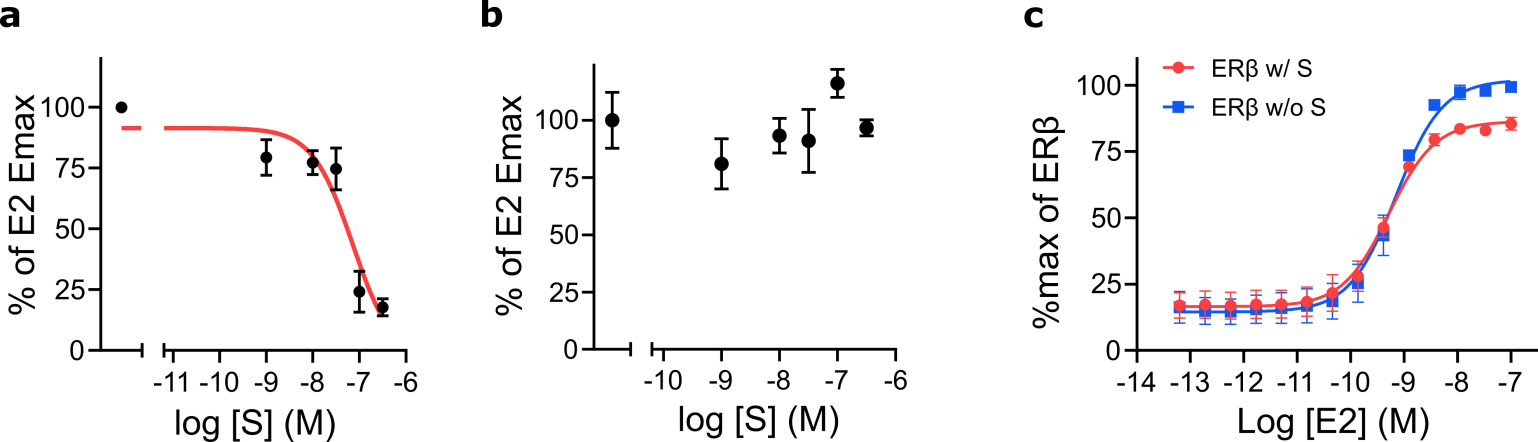

**Extended Data Figure 6.** S trimer (a), but not S-RBD (b), inhibits E2-induced ERα DNA binding in MCF-7 nuclear extracts. Transfected S inhibits E2-stimulated ERβ transcriptional activity (c). Data are shown as mean ± SEM.

**Extended Data Figure 7.**

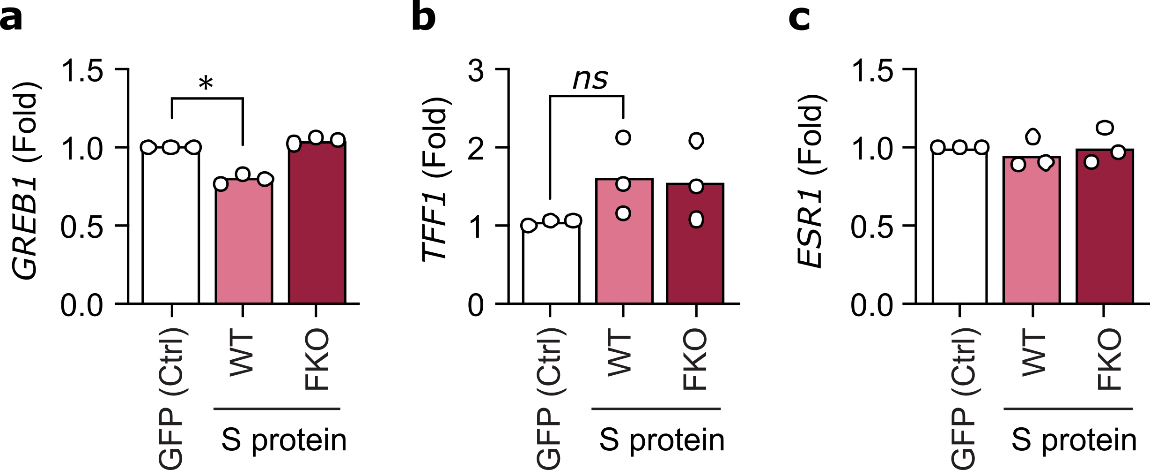

**Extended Data Figure 7.** S overexpression dysregulates ERα target genes. Steady-state levels of the indicated transcripts were compared by qPCR in steroid-deprived MCF7 cells, 48 h after transfection with green-fluorescent protein (GFP), or either wildtype (WT) or the furin cleavage site mutant (FKO) S protein expression vector. Bars represent the mean fold change.

**Extended Data Figure 8.**

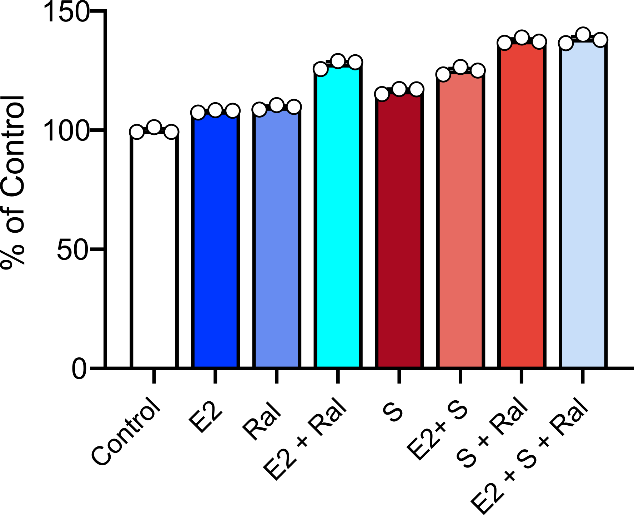

**Extended Data Figure 8.** BrdU Proliferation assay for MDA-MB-231 cell line. All the treatments alone or in combination do not significantly affect cell proliferation. The assay was performed in triplicate. Results are shown as mean ± SD of one representative experiment.

**Extended Data Figure 9.**

**a**

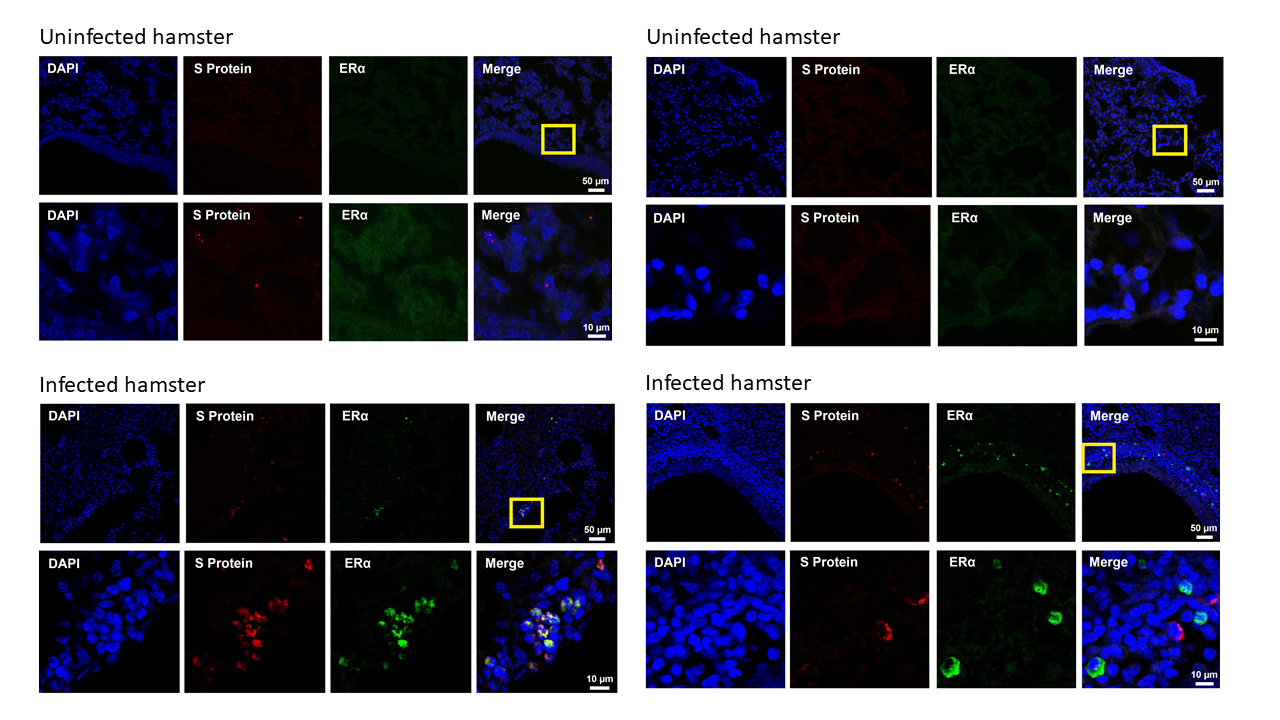
**b**

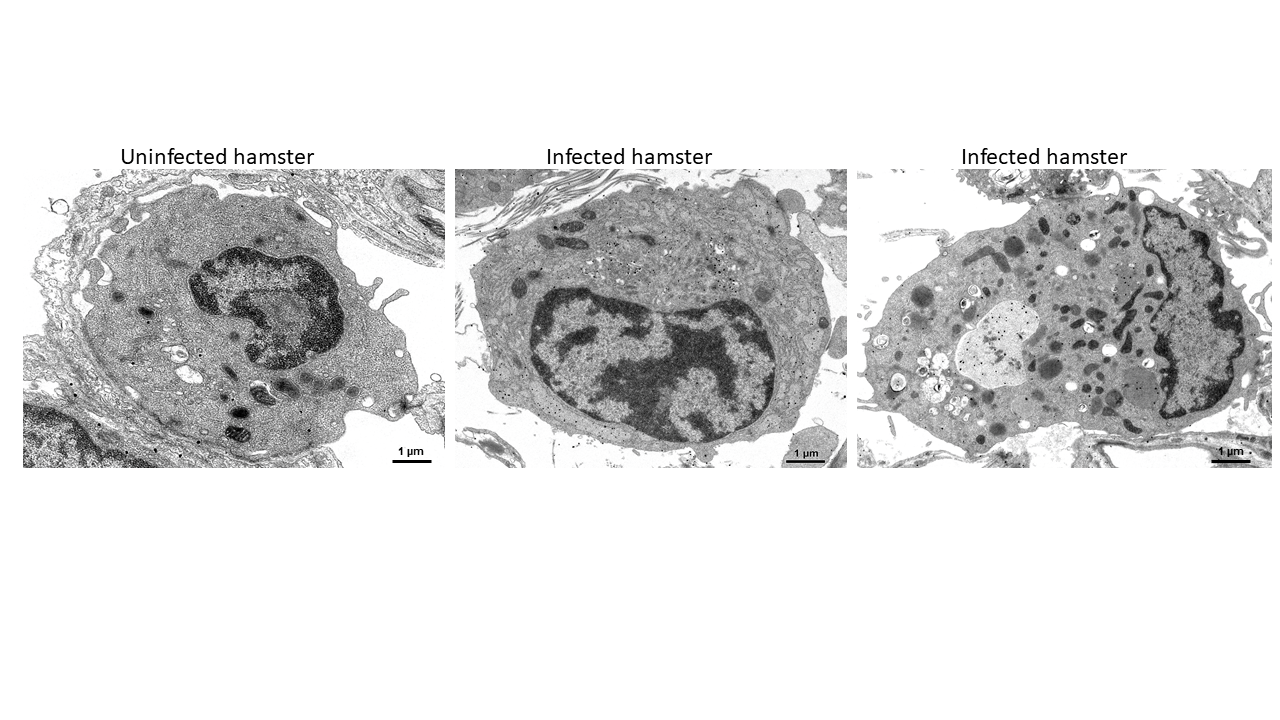
**c**

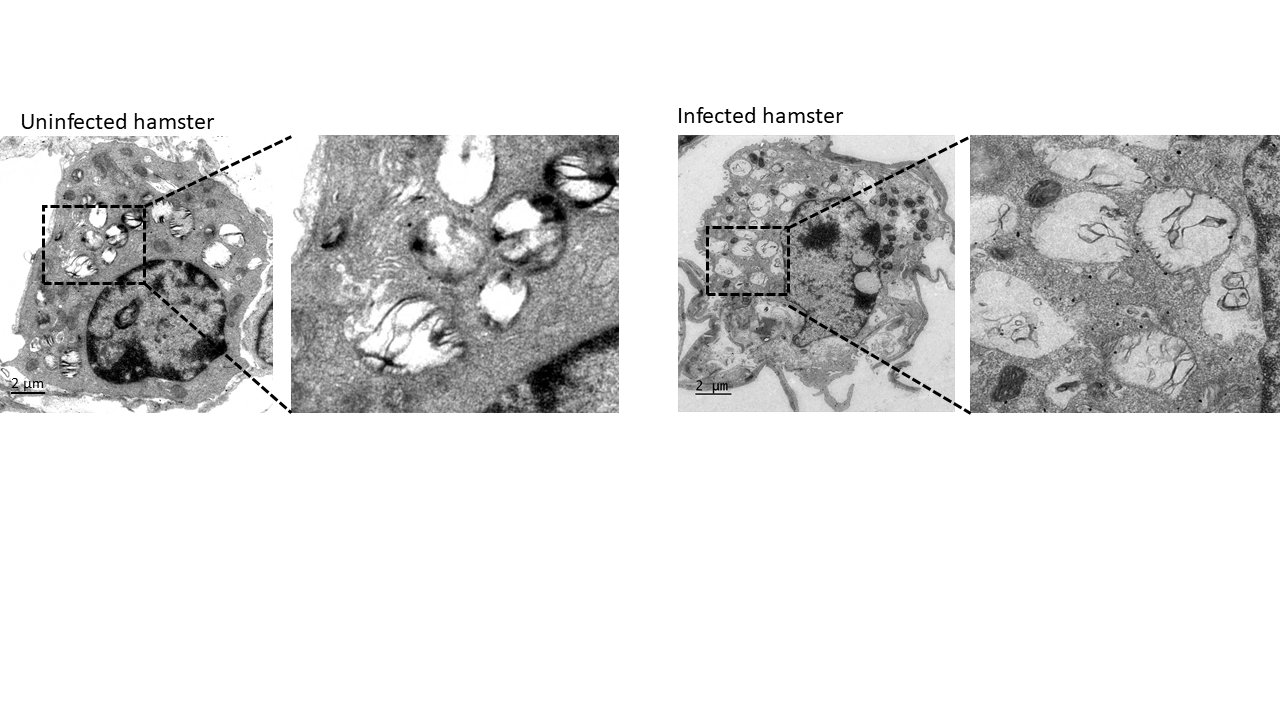

**Extended Data Figure 7.** SARS-CoV-2 infection induces cytoplasmic expression of ERα protein in hamsters. Immunohistochemistry showing colocalization of S and ERα immunoreactivity in infected lung hamster (a). Immunogold EM showing ERα-bound gold nanoparticles in hamster alveolar macrophages (b) and pulmonary type II cells (c).

**Extended Data Figure 10.**

**a**

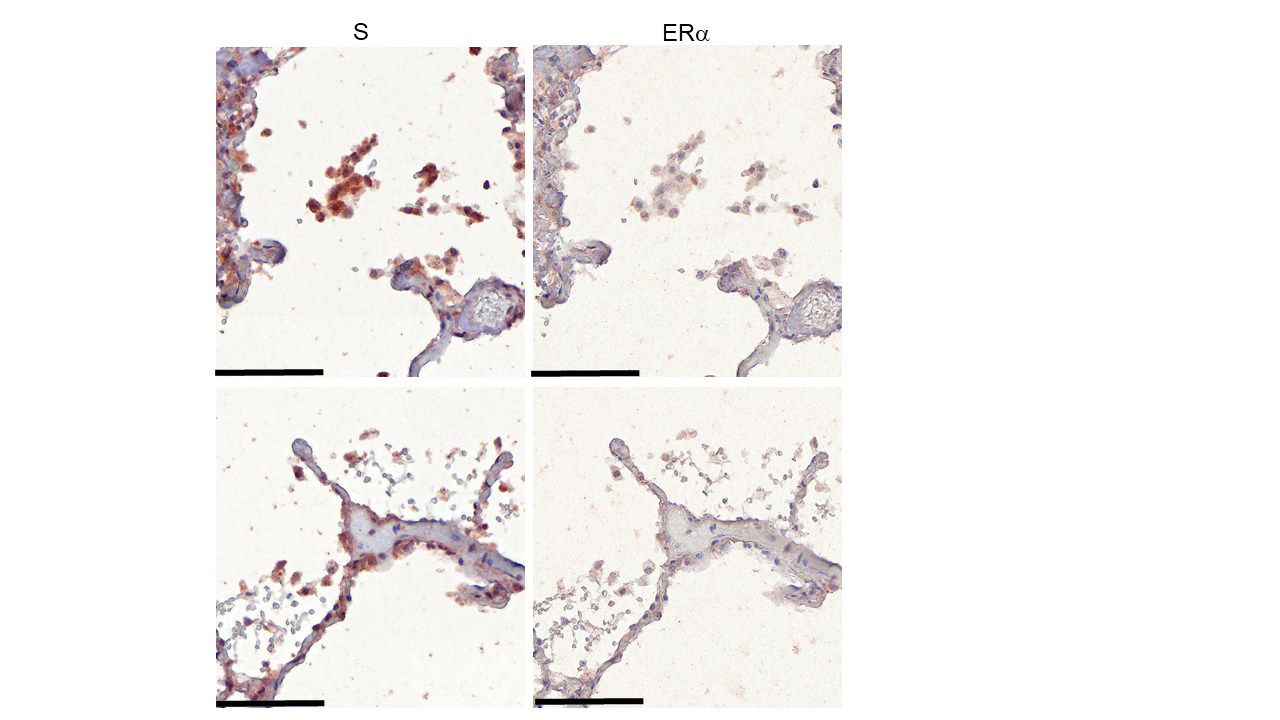

**b**

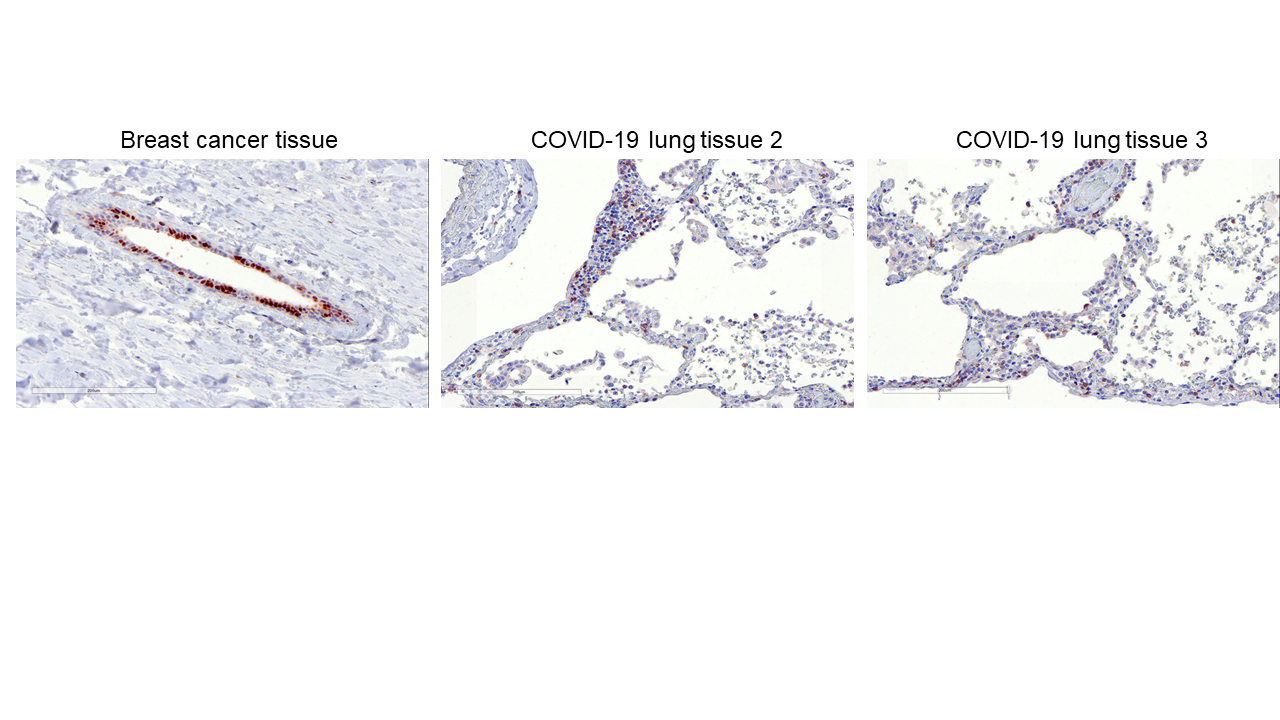

**Extended Data Figure 10.** S and ERα immunostaining in SARS-CoV-2-infected human lung showing S-ERα colocalization. Scale bar 100 nm (a). Immunohistochemistry showing ERα immunoreactivity in breast cancer tissue. Scale bar 200 nm (b).

**Extended Data Figure 11.**

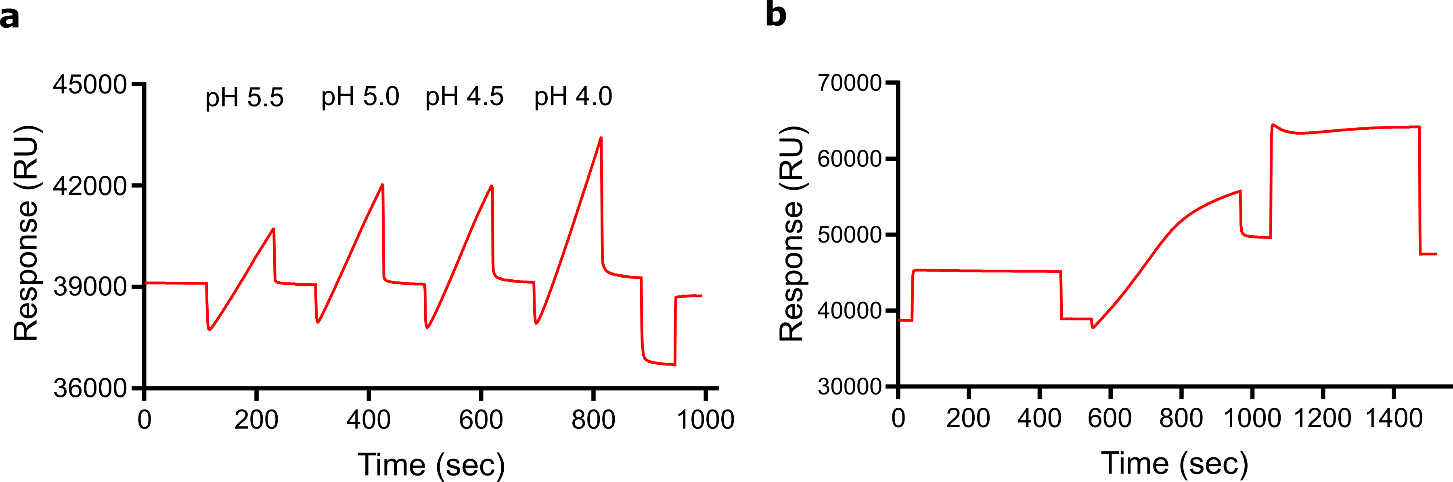

**Extended Data Figure 11.** (a) The sensorgram shows the effect of buffer pH on immobilization of S to the chip surface. (b) Sensorgram illustrating the immobilized amount of S at pH 4.

**Extended Data Table 1**

| Database ID | Mean signal used | Mean Z-Factor | Mean Z-Score | Mean CI P-Value | Mean CV | Description |
| --- | --- | --- | --- | --- | --- | --- |
| NP_001019799.1 | 2553.66 | 0.44708 | 0.829635 | 0.043687 | 0.04742 | NRP1 / Neuropilin-1 Protein |
| BC020803.1 | 6186.558 | 0.57108 | 7.054942 | 0.002179 | 0.09359 | Developmentally regulated GTP binding protein 1 (DRG1) |
| BC033792.1 | 2805.59 | 0.671277 | 2.627567 | 0.013592 | 0.04339 | Tumor protein D52-like 3 (TPD52L3) |
| NM_016059.3 | 3130.755 | 0.641917 | 3.062002 | 0.010238 | 0.024877 | Peptidylprolyl isomerase (cyclophilin)-like 1 (PPIL1) |
| NM_016508.2 | 5234.063 | 0.594467 | 5.65815 | 0.003209 | 0.081497 | Cyclin-dependent kinase-like 3 |
| NM_022140.2 | 2422.185 | 0.554883 | 1.70084 | 0.043508 | 0.058637 | Band 4.1-like protein 4A |
| NM_145010.1 | 8754.735 | 0.656837 | 9.09587 | 0.001407 | 0.074957 | chromosome 10 open reading frame 63 (C10orf63) |

The data was obtained from three independent Protoarrays

**Extended Data Table 1.** Proteins identified by protein-protein interaction with [^125^I]S using the Protoarray platform.

**Extended Data Table 2**

| **INTERACTION PROTEINS** | **SCORE** |
| --- | --- |
| ERα > NCOA1 | 0.997 |
| ERα > NCOA2 | 0.995 |
| ERβ > NCOA1 | 0.994 |
| ERβ > NCOA2 | 0.971 |
| ERβ > NCOA3 | 0.976 |

**Extended Data Table 2.** Scores showing strong interaction between ERα/β and NCOAs.

**Extended Data Table 3.**

| Virus Species | UNIPROT CODE |
| --- | --- |
| Severe acute respiratory syndrome coronavirus 2 (2019-nCoV) (SARS-CoV-2) | SPIKE_SARS2 |
| Severe acute respiratory syndrome coronavirus (SARS-CoV) | SPIKE_SARS |
| Middle East respiratory syndrome-related coronavirus (MERS-CoV) | SPIKE_MERS1 |
| Human coronavirus HKU1 (isolate N5) (HCoV-HKU1) | SPIKE_CVHN5 |
| Human coronavirus OC43 (HCoV-OC43) | SPIKE_CVHOC |
| Murine coronavirus (strain A59) (MHV-A59) (Murine hepatitis virus) | SPIKE_CVMA5 |
| Murine coronavirus (strain 4) (MHV-4) (Murine hepatitis virus) | SPIKE_CVM4 |
| Murine coronavirus (strain JHM) (MHV-JHM) (Murine hepatitis virus) | SPIKE_CVMJH |
| Murine coronavirus (strain JHMV / variant CL-2) (MHV) (Murine hepatitis virus) | SPIKE_CVMJC |

**Extended Data Table 3.** Coronavirus’ S proteins used for sequence and amino acids pattern analysis.
